# Discovery of multi-state gene cluster switches determining the adaptive mitochondrial and metabolic landscape of breast cancer

**DOI:** 10.1101/2023.02.17.528922

**Authors:** Michela Menegollo, Robert B. Bentham, Tiago Henriques, Seow Qi Ng, Ziyu Ren, Clarinde Esculier, Sia Agarwal, Emily Tong, Clement Lo, Sanjana Ilangovan, Zorka Szabadkai, Matteo Suman, Neill Patani, Avinash Ghanate, Kevin Bryson, Robert C. Stein, Mariia Yuneva, Gyorgy Szabadkai

**Author notes:** **Additional information:**. shared first authorship (Michela Menegollo, Robert B. Bentham). shared senior authorship (Mariia Yuneva, Gyorgy Szabadkai). RB: Cancer Genome Evolution Research Group, Cancer Research UK Lung Cancer Centre of Excellence, University College London Cancer Institute, London, WC1E 6BT, UK; KB: University of Glasgow, School of Computing Science, Glasgow G12 8RZ, UK. correspondence to: Gyorgy Szabadkai, Michela Menegollo, Robert B. Bentham.

## Abstract

Adaptive metabolic switches are proposed to underlie conversions between cellular states during normal development as well as in cancer evolution, where they represent important therapeutic targets. However, the full spectrum, characteristics and regulation of existing metabolic switches are unknown. We propose that metabolic switches can be recognised by locating large alternating gene expression patterns and associate them with specific metabolic states. We developed a method to identify interspersed genesets by massive correlated biclustering (MCbiclust) and to predict their metabolic wiring. Testing the method on major breast cancer transcriptome datasets we discovered a series of gene sets with switch-like behaviour, predicting mitochondrial content, activity and central carbon fluxes in tumours associated with different switch positions. The predictions were experimentally validated by bioenergetic profiling and metabolic flux analysis of ^13^C-labelled substrates and were ultimately extended by geneset analysis to link metabolic alterations to cellular states, thus predicting tumour pathology, prognosis and chemosensitivity. The method is applicable to any large and heterogeneous transcriptome dataset to discover metabolic and associated pathophysiological states.

**Statement of significance:** We present a novel method to identify the transcriptomic signatures of metabolic switches underlying divergent routes of cellular transformation. We illustrate the power of the method by stratifying breast cancer into metabolic subtypes, predicting their biology, architecture and clinical outcome.

## Introduction

Cancer development and evolution entail diverging routes of adaptation in cellular energetics and metabolism to oncogenic signals and the tumour environment^1–3^. These adaptive metabolic rearrangements often result in tumour specific vulnerabilities to pharmacological or dietary targeting, or to the contrary, promote therapy resistance^4^. Evaluating patterns of tumour metabolic adaptation is therefore essential to understanding cancer biology and will allow development of novel therapeutic strategies.

No comprehensive evaluation of metabolic adaptation patterns in cancer has yet been completed. Adaptation patterns are presumed to accompany both transient alterations of cellular phenotype and the inheritable phenotype switches that underlie transitions in cellular states. States can be determined by cell type, their environment and by cellular position in the differentiation tree. Transitions in cell state would therefore follow changes in differentiation status and transdifferentiation, including epithelial-mesenchymal transition (EMT), cellular senescence, or reversion to an immature stem or progenitor phenotype^5,6^, and to other cell type specific state transitions^7^. There has been recent interest in understanding metabolic adaptations during cell state transitions^8,9^ leading to a considerable body of knowledge of cell state specific metabolic alterations at the metabolite and flux levels^2,10–14^. Using metabolites and fluxes as exclusive readouts however may not address the question of whether the altered metabolic states represent *bona fide* functional adaptations or are simply mass action driven rearrangements of metabolic networks^15^. Moreover, metabolic measurements themselves are insufficient to allow exploration of the relationship and the co-regulation of the functional adaptation with cellular state.

Gene expression pattern (GEP) alterations almost invariably accompany cellular state transitions and transformation and bear important prognostic value^16–18^. Accordingly, GEPs can provide information that links cellular state to metabolic adaptation. This can be evaluated by exploring the metabolic transcriptome in large scale GEPs, including expression of metabolic pathway components and nuclear encoded mitochondrial genes which together represent >15% of the genome. Large patterns of alteration in the transcriptome, often involving >1000 genes, can arise either from genetic (oncogenic mutations) or non-genetic (environmental) influences. These ultimately determine cellular phenotype and the associated cellular metabolic network^19^. Earlier studies on cancer related metabolic rearrangements mostly focused on the effect of cell-autonomous oncogenic signals on specific metabolic pathways^20,21^, thereby delineating either the metabolic switches required for increased proliferation and growth or adaptation to hypoxia^1,22^. However, it is now well recognised that other cellular states or features can also be associated with altered gene expression patterns affecting metabolic networks, including not only the fundamental EMT transdifferentiation in cancer metastasis, but also other pathological and physiological states^16,17,23,24^. In order to find patterns of metabolic adaptation, we hypothesised that alterations in metabolic gene expression patterns reflect *bona fide* adaptive transcriptional regulation, driven by a specific cell state. Accordingly, metabolic switches follow transitions between cell states, and apart from being clear pharmacological targets^25^, may well be used for important pathological predictions associated with cell states. To test this hypothesis, here we aimed to identify differential regulation patterns in the full metabolic transcriptome and link these quantitatively with the large scale GEPs associated with different cancer cell states. Since the transcriptome does not scale linearly with the proteome and the metabolome^26^, we aimed to correlate these states with metabolic and energy flux measurements experimentally. Finally, we applied statistical methods to understanding the correlation of GEPs with tumour biology and establishing their prognostic value. Identifying these patterns will help the unbiased mapping of potential metabolic adaptation strategies in tumours, understanding and exploitation of metabolic vulnerabilities in different cell states represented as tumour subtypes, in particular to recognition of and targeting adaptive patterns resulting in drug-resistant states.

Few previous studies analysed large scale metabolic gene expression data across multiple cancer types, mostly revealing widespread deregulation of metabolic pathways compared to adjacent normal tissue. These studies used either diverse pathway scoring methods based on differential gene expression in numerous cancer types and/or subtypes^16,18,27,28^, or analysed genomic changes associated with transcriptional adaptation to the environment^22^. For example, in order to investigate shared deregulation of individual metabolic pathways genes, Reznik et al.^17^ analysed differential gene co-expression in a number of cancer types, revealing recurrent loss of co-expression of a set of genes, including mitochondrial electron transport chain components. However, these studies did not account for large, correlated expression patterns in broader sets of genes, which inherently underlie the complete metabolic rewiring of phenotypic (cell state) switching. Such patterns are likely to be aligned with specific cell states and could reveal subtype specific dependencies that provide novel pharmacological targets or predict pathological features and clinical behaviour. In addition, defining these patterns will help answer questions such as what drives the deregulation of metabolic pathways, and whether it is adaptive and thus targetable, as well as how gene expression and metabolic fluxes correlate. No current approaches to the best of our knowledge have addressed this problem.

To undertake the unbiased discovery of large co-regulated gene expression patterns, we have simplified our model to include the concerted on and off switching of large gene sets^29^, such as bi- and multistate, or bi- and multistable gene switches. These switches thus comprise anticorrelated gene sets in a selected set of samples representing two different cell states or cancer subtypes. We have created a pipeline for the discovery and further analysis of these large gene expression switches which can be used as a resource to analyse any large heterogeneous dataset. In order to demonstrate the workflow and the value of the resource, we describe here its use in the discovery of metabolic and mitochondrial states (subtypes) of breast cancer, aligning these with cell state determined by the cell of origin. Firstly by using the central tool of the resource, MCbiclust^30^ applied to a number of large breast cancer transcriptome datasets, we have discovered three massive anticorrelated gene sets. These biclusters define breast cancer metabolic subtypes which are dominated by mitochondrial genes and are likely the result of metabolic switches that follow the cell of origin pattern. Next, we correlated the gene expression patterns with metabolic fluxes, measured in cellular models by mass spectrometry using heavy carbon labelled substrates. Finally, we have demonstrated the utility of the resource through its ability to predict (i) the metabolic phenotype of cancer cell lines and cancer tissue; (ii) the metabolic wiring associated with certain cell states and phenotypes; and (iii) clinical outcomes of a subset of estrogen receptor (ER) positive tumours which offers an opportunity to modify current treatment algorithms.

## Materials and Methods

### Resources

| REAGENT or RESOURCE | SOURCE | IDENTIFIER |
| --- | --- | --- |
| Antibodies |  |  |
| Rabbit monoclonal anti-Hexokinase 1 | Cell Signaling Technology | Cat# 2024, RRID:AB_2116996 |
| Rabbit monoclonal anti-Hexokinase 2 | Cell Signaling Technology | Cat# 2867, RRID:AB_2232946 |
| Rabbit polyclonal anti-Triosephosphate isomerase –N-terminal | Abcam | Cat# ab135532 |
| Monoclonal antibody anti-Glyceraldehyde-3-phosphate dehydrogenase | Millipore | Cat# MAB374, RRID:AB_2107445 |
| Rabbit monoclonal anti-Pyruvate kinase M1 isoform | Cell Signaling Technology | Cat# 7067, RRID:AB_2715534 |
| Rabbit monoclonal anti-Pyruvate kinase M2 isoform | Cell Signaling Technology | Cat# 4053, RRID:AB_1904096 |
| Rabbit monoclonal anti-Lactate dehydrogenase A | Cell Signaling Technology | Cat# 3582, RRID:AB_2066887 |
| Rabbit monoclonal anti-Lactate dehydrogenase B | Abcam | Cat# ab53292, RRID:AB_2234531 |
| Rabbit polyclonal anti-BRP44L (MPC1) | Abcam | Cat# ab74871, RRID:AB_1523226 |
| Rabbit monoclonal anti-MPC2 | Cell Signaling Technology | Cat# 46141, RRID:AB_2799295 |
| Mouse monoclonal anti-Pyruvate dehydrogenase E1-alpha subunit | Abcam | Cat# ab110330, RRID:AB_10858459 |
| Rabbit polyclonal anti-pyruvate dehydrogenase E1-alpha subunit (phospho S293) | Abcam | Cat# ab92696, RRID:AB_10711672 |
| Rabbit polyclonal anti-IDH1 | Proteintech Group | Cat# 12332-1-AP,<br>RRID:AB_2123159 |
| Mouse monoclonal anti-IDH2 | Abcam | Cat# ab55271,<br>RRID:AB_943793 |
| Rabbit polyclonal anti-IDH3A | Abcam | Cat# ab58641,<br>RRID:AB_2295799 |
| Rabbit polyclonal<br>anti-glutaminase | Abcam | Cat# ab93434,<br>RRID:AB_10561964 |
| Rabbit monoclonal anti-GLUD1 | Abcam | Cat# ab166618,<br>RRID:AB_2815030 |
| Rabbit polyclonal anti-GOT1 | Proteintech Group | Cat# 14886-1-AP,<br>RRID:AB_2113630 |
| Rabbit polyclonal anti-GOT2 | Proteintech Group | Cat# 14800-1-AP,<br>RRID:AB_2247898 |
| Rabbit polyclonal anti-ME1 | Abcam | Cat# ab97445,<br>RRID:AB_10679994 |
| Rabbit monoclonal anti-ME2 | Abcam | Cat# Ab139686 |
| Rabbit monoclonal anti-PCB | Abcam | Cat# ab128952,<br>RRID:AB_11144490 |
| Rabbit polyclonal anti- 3PGDH<br>(PHGDH) | Santa Cruz Biotechnology | Cat# sc292792 |
| Mouse monoclonal anti-PPAT | Abcam | Cat# ab128116,<br>RRID:AB_11143951 |
| Rabbit polyclonal anti-PSAT1 | Proteintech Group | Cat# 20180-1-AP,<br>RRID:AB_10665948 |
| Rabbit monoclonal anti-MDH1 | Abcam | Cat# ab180152 |
| Mouse anti - GLUL | BD Biosciences | Cat# 610518<br>RRID:AB_397880 |
| Rabbit monoclonal anti-GFPT1<br>[EPR4854] | Abcam | Cat# ab125069,<br>RRID:AB_10975709 |
| Recombinant anti-GFPT2 | Abcam | Cat# ab190966,<br>RRID:AB_2868470 |
| Total OXPHOS rodent antibody cocktail | Abcam | Cat# ab110413, RRID:AB_2629281 |
| Mouse monoclonal anti-Actin | Santa Cruz Biotechnology | Cat# sc-56459, RRID:AB_830981 |
| Rabbit polyclonal anti-beta tubulin | Santa Cruz Biotechnology | Cat# sc-9104, RRID:AB_2241191 |
| Rabbit polyclonal anti-grp75 | Santa Cruz Biotechnology | Cat# sc-13967, RRID:AB_647720 |
| Chemicals, Peptides, and Recombinant Proteins |  |  |
| [U-<sup>13</sup>C] glucose | Cambridge Isotope Laboratories, Inc. | CLM-1396 |
| [U-<sup>13</sup>C] glutamine | Cambridge Isotope Laboratories, Inc. | CLM-1822 |
| [U-<sup>13</sup>C] pyruvate | Cambridge Isotope Laboratories, Inc. | CLM-1575 |
| Pyruvate | Thermo Fisher Scientific | Cat# 11360070 |
| Glutamine | Thermo Fisher Scientific | Cat# 25030081 |
| Glucose | Thermo Fisher Scientific | Cat# A2494001 |
| Normocin | InvivoGen | ant-nr-1 |
| Hoechst 33342 | Thermo Fisher Scientific | Cat# 62249 |
| Tetramethylrhodamine, Methyl Ester, Perchlorate (TMRM) | Thermo Fisher Scientific | Cat# T668 |
| Quant-iT™ PicoGreen™ dsDNA Assay Kits and dsDNA Reagents | Thermo Fisher Scientific | Cat# P11495 |
| Rotenone | Sigma-Aldrich | Cat# R8875 |
| Carbonyl cyanide 4-(trifluoromethoxy)phenylhydraz one (FCCP) | Sigma-Aldrich | Cat# C2920 |
| Antimycin A from Streptomyces sp. | Sigma-Aldrich | Cat# A8674 |
| Oligomycin A | Sigma-Aldrich | Cat# 75351 |
| DMEM, high glucose, pyruvate | Thermo Fisher Scientific (Gibco) | Cat# 41966029 |
| DMEM, high glucose, glutaMAX supplement, pyruvate | Thermo Fisher Scientific (Gibco) | Cat# 31966021 |
| DMEM, no glucose, no glutamine, no phenol red | Thermo Fisher Scientific (Gibco) | Cat# A1443001 |
| Critical Commercial Assays |  |  |
| Glutamine/Glutamate-Glo Assay | Promega | Cat# J8022 |
| CellTiter-Glo Luminescent Cell Viability Assay | Promega | Cat# G7570 |
| Seahorse XF Cell Mito Stress Test Kit | Agilent | Cat# 103015-100 |
| Experimental Models: Cell Lines |  |  |
| MCF7 | ATCC | HTB-22 |
| T47D | ATCC | HTB-133 |
| Hs578T | ATCC | HTB-126 |
| MDA-MB436 | ATCC | HTB-130 |
| HCC1143 | ATCC | CRL-2321 |
| Source Data |  |  |
| METABRIC | Synapse: syn1757063 | https://doi.org/10.7303/syn1688369 |
| TCGA | NCBI dbGaP | phs000178.v11.p8 |
| OsloVal | Synapse: syn1710395 | https://doi.org/10.7303/syn1688370 |
| Software and Algorithms |  |  |
| R v4.2.0 | R Foundation for Statistical Computing | https://www.r-project.org/ |
| MCbiclust 1.22.0 | Bentham et al.<sup>30</sup> | https://doi.org/doi:10.18129/B9.bioc.MCbiclust |
| ComplexHeatmap | Gu<sup>79</sup> | https://doi.org/doi:10.18129/B9.bioc.ComplexHeatmap |
| GSVA 1.46.0 | Hänzelmann et al<sup>44</sup> | https://doi.org/doi:10.18129/B9.bioc.GSVA |
| g:Profiler | Raudvere et al<sup>80</sup> | http://biit.cs.ut.ee/gprofiler/gost |
| Fiji | Schindelin et al.<sup>81</sup> | https://imagej.net/software/fiji/ |
| Prism 9.0 | GraphPad Software | https://www.graphpad.com/ |
| Other |  |  |
| SnakeSkin Dialysis Tubing, 3.5K MWCO, 35 mm dry I.D., 35 feet | Thermo Fisher Scientific | Cat# 88244 |
| Seahorse XF96 cell culture microplates | Agilent | Cat# 102416 |
| Falcon® 96-well Black/Clear Flat Bottom TC-treated Imaging Microplate | Corning Life Sciences | Cat# 353219 |

### Resource availability

#### Lead contact

Further information and requests for resources and reagents should be directed to and will be fulfilled by the lead contact, Gyorgy Szabadkai.

#### Materials availability

This study did not generate new unique reagents.

#### Data and code availability

All data are available via links provided in the Methods section and Key Resources table. Code for the full analysis pipeline is available from the authors upon request.

### Experimental model and subject details

#### Cell lines

Human breast cancer cell lines MCF7, T47D, Hs578t, MDA436 and HCC1143 were purchased from ATCC. Their identity was confirmed through cell line authentication carried out by BMR Genomics (Italy) by amplification of 23 STR loci (PowerPlex Fusion System kit, Promega). Cells were cultured in Dulbecco’s modified Eagle’s medium (DMEM, #41966029), supplemented with 10% Fetal Bovine Serum (FBS) (Thermo Fisher Scientific) and Normocin (0.1 mg/mL, InvivoGen) and maintained in a 37°C incubator set with 5% CO2 and 95% humidity. Cell numbers for each experimental setting were counted using a hemocytometer. For the restricted nutrient condition and stable isotope labelling experiments, cells were cultured using DMEM (no glucose, no glutamine, no phenol red, Thermo Fisher Scientific #A1443001) complemented with glucose, glutamine and pyruvate, as indicated. For the stable isotope labelling, the required amount of labelled compounds, [U-^13^C] glucose, [U-^13^C] glutamine and [U-^13^C] pyruvate, were added. FBS was dialyzed using a SnakeSkin dialysis tubing (Thermo Fisher Scientific) and filtered using 0.22 µm PES filter before being added in 10% proportion, together with Normocin. The conditions were named according to their nutrient concentrations either stably labelled or not (Figure 4A): high and low glutamine (H/L = 1mM/0.1mM) levels were combined with the presence and absence of pyruvate (+/- = 1mM/0mM), glucose was kept at 10 mM.

### Methods details

#### Discovery of large gene cluster switches with MCbiclust

MCbiclust (doi:10.18129/B9.bioc.MCbiclust, current version 1.2.1, R package available in Bioconductor^82^) was used to identify gene cluster switches as described previously^30,31^ on the METABRIC^33^, TCGA^34^ (BRCA) and Oslo2^35^ datasets. We have used two defined and two random starting genesets of approximately 1000 genes. The defined genesets were either only mitochondrial (Mitocarta 1.0^83^) or mitochondria related, by choosing the 1000 genes most correlated with the mitochondrial ribosomal gene MRPL58 across the METABRIC dataset. MCbiclust was run for 1000 iterations using each geneset on the University College London high performance cluster. The initial bicluster seeds found are shown in Figure S1, following the selection of the highest correlating gene groups using the HclustGenesHiCor function. The silhouette method was used to determine the number of independent biclusters. Biclusters were then extended (i) by calculating the correlation vector (CV) values for the whole transcriptome using the CVEval function, and (ii) to all samples using SampleSort. The correlation pattern was summarised using principal component analysis (PCA) of the CV values using PC1VecFun. The bifurcating patterns, defining the forks (switch positions), were plotted using the sort order (index) and PC1 values for each sample (see Figures 1B, S1).

**Figure 1.**
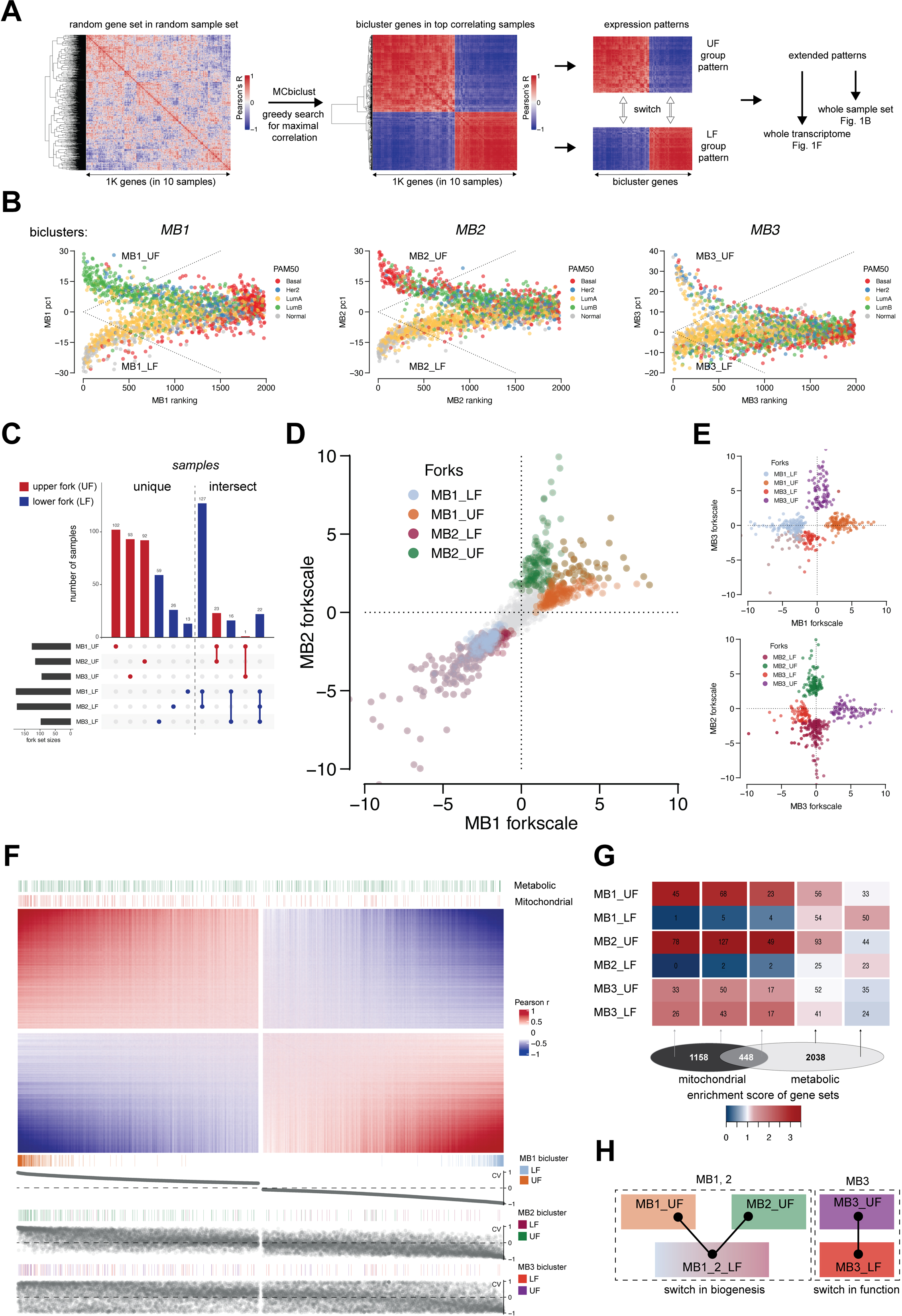
Identification of large scale transcriptome switches. **A.** Geneset switch identification in bicluster samples. MCbiclust starts with a random geneset in a random sample set, and performs a greedy search aiming for the sample set across which the geneset has the highest correlation. Since the *absolute* value of correlation is maximised during the search, the resulting top correlating samples will display two genesets with sharp anticorrelation (switch) between them. The genesets and samplesets can be extended to the whole transcriptome and sampleset by quantifying correlations (see main text), which provides the basis for gene set enrichment analysis **B.** Visualisation of individual switches (MB_1, MB_2, MB_3) in 2D distribution by PC1 and ranking index in the METABRIC dataset. PAM50 classifiers are overlaid to show the differential distribution of samples in the different switches (biclusters). Dashed lines represent the thresholds for belonging to a fork, calculated as 0.04 * forkscale (see Methods). **C.** UpSet matrix and plots of aggregates of sample intersections between upper and lower positions of each switch (MB_1, MB_2, MB_3). Unique and overlapping (intersect) samples are shown separately. **D, E.** Interswitch relationships are shown in 2D distribution plots of samples by two biclusters (switches). The axes represent the scale between the upper (UF) and lower (LF) of each bicluster. The distribution of samples according to their MB_1 vs MB_2 forkscale value (See Methods) is shown in **D**, whilst the combinations of MB_3 with MB_1 or MB_2 are shown in **E**. **F.** Main heatmap shows distribution of the correlation values (CVs, Pearson r) of nearly the whole transcriptome in the MB_1 bicluster; a small set of midrange values were hidden (see gaps in heatmap). Annotations show the locations of mitochondrial and metabolic genes (top), and that of the bicluster genes (bottom). For comparison between the switches, bottom annotation curves show the CV values of each sample in the three biclusters (switches). **G.** Fold enrichment of mitochondrial and metabolic (with intersections) genes in the anticorrelated gene sets. Heatmap shows enrichment values of each geneset in each position of all switches. Geneset intersections are illustrated at the bottom. **H.** Scheme of switches. MB1 and MB2 share their LF, thus representing a multistate switch, while MB3 is an independent bistable switch, both MB1_2 and MB_3 representing different switch mechanisms (see Main text).

#### Gene set enrichment and pathway analysis

Functional analysis of gene clusters were performed with g:Profiler^80^. The default KEGG data source was used in g:GOSt for functional profiling of all metabolic pathways. The KEGG results were manually filtered to include only metabolic pathways. For mitochondrial functional profiling the custom MitoCarta 3.0 orthology was used as data source^36^ in g:Profiler. Signalling pathway activity was assessed using the PARADIGM integrated pathway levels (IPLs) from the PARADIGM^39^ analysis on a pan-cancer dataset^37^ (https://www.synapse.org/#!Synapse:syn5633407), filtered for BRCA samples. Gene set variation analysis for molecular signature identification was performed using the GSVA 1.46.0 package^44^.

#### Assessment of mitochondrial activity

Oxygen consumption rate was measured with the Seahorse XFe96 bioanalyzer using the Seahorse XF Cell Mito Stress Test Kit (Agilent), or using High-Resolution Respirometry (Oxygraph-2K, Oroboros Instruments, Austria). For the Seahorse measurements cells were seeded on XF96 cell culture microplates (Agilent) 2 days before the experiment (1 × 10^4^ cells/well). For the experiment, the culture medium was replaced with Seahorse XF Base medium (Agilent) supplemented with pyruvate, glutamine and glucose (Thermo Fisher Scientific) as indicated, and incubated for 30 min at 37 °C in a CO_2_-free incubator before loading into the Seahorse Analyser. After measuring basal respiration, the drugs oligomycin A (5 µM), FCCP (1 µM), and rotenone/antimycin A (0.5 µM/0.5 µM) were added to each well in sequential order. Data were analysed using the XF Cell Mito Stress Test Report Generator. After the assay, cells were stained with Hoechst 33342 (1 µg/mL; Thermo Fisher Scientific) for 30 min. ImageXpress Micro XL was then used for cell nuclei (cell numbers) counts in each well.

For High-Resolution Respirometry the Oxygraph-2K instrument was used. Prior to the assay, OROBOROS chambers were calibrated with the recording media (DMEM #A1443001, glucose (10 mM), glutamine (1mM), pyruvate (1mM), HEPES (10 mM)). Confluent 10 cm plates of cells were trypsinized and re-suspended in the same media. 2.5*10^6^ cells were added to each chamber, and the O_2_ flow signal was allowed to stabilise to basal respiration rate. Drugs were added to chambers using in the following concentrations and order: Oligomycin (2.5µM), FCCP (2µM) and Antimycin A (2.5 µM). Respiratory rate given as oxygen flow in pmol/min/cell was recorded. Data collection and analysis was performed using Datlab 5.0 software.

#### Imaging

For imaging overall mitochondrial structure and function in a large cell population, a wide-field high content imaging system (ImageXpress MicroXL, Molecular Devices) was used. Cells were seeded in 96-well black clear thin bottom tissue culture treated imaging plates (Corning). For mitochondrial membrane potential measurements cells were incubated with 30nM TMRM (Thermo Fisher Scientific) in recording media (DMEM #A1443001, glucose (10 mM), glutamine (1mM), pyruvate (1mM), HEPES (10 mM)) for 30 minutes prior to starting imaging and left for the whole time of the experiment at 37°C. For labelling the nuclei cells were co-loaded with Hoechst 33342 (1 µg/mL; Thermo Fisher Scientific) for 30 min. For TMRM intensity quantification a 20X Nikon (S PLAN FLUOR ELWD 0.45 NA) air objective was used. Images were analysed with the integrated metaXpress imaging and analysis software using the granularity analysis module. Following local background subtraction, the average TMRM intensity/field was used as readout.

For morphology analysis a Nikon 40X (Plan Apo 0.95 NA) air objective was used. Cells were co-loaded with TMRM (30 nM), picoGreen (2.5µL/mL) and Hoechst 33342 (1 µg/mL; Thermo Fisher Scientific) in recording media (DMEM #A1443001, glucose (10 mM), glutamine (1mM), pyruvate (1mM), HEPES (10 mM)) for 30 minutes prior to starting imaging. TMRM was left for the whole time of the experiment at 37°C. Cells were imaged using a custom protocol to image 3 wavelengths (Lumencor solid state illumination with Semrock [Brightline] filters (nm) ex377/50 em447/60 for Hoechst 3342; ex472/30 em520/35 nm for picoGreen, ex531/40 em593/40 for TMRM), 16 fields/well with a Nikon 40X Plan Apo NA 0.95 air objective, binning 1 with a CMOS detector. TMRM and picoGreen Images were analysed with MetaXpress 5.0 software, using the granularity module, total mitochondrial area and average object size as readout from the TMRM images. For picoGreen analysis, the integrated pit intensity (granularity module) was used as a readout following ratioing with the Hoechst images, background subtraction and thresholding.

#### Determination of total cellular ATP

Total cellular ATP concentration was measured using CellTiter-Glo® Luminescent Cell Viability Assay (Promega). The method determines the number of viable cells based on the quantification of the ATP present, as readout of metabolically active cells, thus allowing the estimation of total cellular ATP produced by a known number of seeded cells. In detail, 5K cells were seeded in an opaque black 96-well plate and treated with the experimental media (Figure 4A) for 24h. Wells containing the experimental medium only (the blank - for removing the medium background) and 10nM ATP standard (on which the samples luminescence was normalised to) were included. The protocol was carried out according to the manufacturer’s instructions. Briefly, i): plates were equilibrated at room temperature (RT) for approximately 30 minutes, avoiding temperature gradients which could cause uneven signals; ii) a volume of CellTiter-Glo® Reagent equal to the volume of cell culture medium present in each well (e.g., 100μl of reagent to 100μl of medium containing cells for a 96-well plate) was added; iii) 2 minutes of orbital shaking favouring cell lysis; then, iv) plates were left at RT for 10 minutes to stabilise luminescent signal and v) the luminescence was recorded using PerkinElmer EnVision plate reader. For each plate the triplicate values for the samples total ATP produced, the ATP standard and five replicates of medium luminescence for the background were acquired. The average of the replicates was calculated, and the medium background was subtracted. The resulting values were normalised to the ATP standard previously adjusted for the background and further normalised to the protein content, evaluated using the BCA protein assay (Pierce).

#### Western Blotting

For the baseline expression of metabolic enzymes and mitochondrial respiratory chain subunits at protein level, proteins were extracted from cells plated in 10 cm dish and cultured in DMEM. Briefly, cells were detached by scraping in cold PBS 1X and pulled down by centrifugation. Whole cell lysate was obtained incubating cell pellet with cold RIPA buffer (150 mM NaCl, 50 mM Tris, 0.5% sodium deossicolate, 0.1% (w/v) SDS, 1% (v/v) Triton) in presence of protease inhibitor cOmplete (Roche), phosphatase inhibitors PhosStop (Roche) and PMSF (Sigma) for 30 minutes on ice and spun down at 4 °C. Protein quantification was done using a BCA protein assay kit (Pierce). Samples were denatured in presence of reducing agents (DTT) at 95°C for 5 minutes, while for probing mitochondrial respiratory chain subunits the optimal temperature was 60°C for 10 minutes. 20 µg of protein lysate was loaded onto a NuPAGE 4-12% bis-tris gel (Thermo Fisher) and run using MOPS 1X buffer. Blotting was done for 2 h at 30 V in a wet system (Invitrogen - Thermo Fisher), by transferring proteins onto nitrocellulose membrane for metabolic enzymes and PVDF for mitochondrial subunits. Membranes were incubated in 5% non-fat dry milk in TBS 1X – 0.1% Tween 20 or 5% BSA in TBS 1X – 0.1% Tween 20 as blocking buffers (according to each antibody datasheet) for 1 h at room temperature. Primary antibody was incubated in blocking buffer overnight at 4°C. The working dilution was 1:1000 for the most of antibodies except for MitoProfile (1:2000), grp75 (1:3000), actin (1:3000). Horseradish-conjugated secondary antibodies (BioRad) were diluted 1:5000 in blocking buffer and membranes were incubated 1 h at room temperature. The blots were visualised using SuperSignal West Pico Chemiluminescent Substrate (Thermo Fisher) and UVITEC Cambridge Mini HD9 Imaging System (Eppendorf). If necessary, membranes were stripped, blocked and probed again with a different primary antibody.

#### Measurement of glutamine and glutamate concentration in the media

The concentration of glutamine and glutamate in the media were measured using the luminescence-based approach Glutamine/Glutamate-Glo TM Assay kit (Promega). The method is based on two steps: i) glutamine is first converted to glutamate by glutaminase; ii) glutamate is oxidised and NADH produced is used for luminescence reading. Briefly, 10k cells were seeded in an opaque black 96-well plate. On the following day, after one wash with warm PBS1X, cells were treated with the experimental media (Figure 4A) for 24 h. Wells containing the experimental medium only (no cells), 5 points glutamate standard curve (to extrapolate glutamine and glutamate concentration) and wells containing PBS1X (as negative control) were included. Wells with only medium represent a positive control and were used as reference of initial glutamine and glutamate concentration in the medium, on which values subtract the glutamine and glutamate concentration obtained from cells containing wells, thus calculating the glutamine uptake and glutamate secreted by the cells. The addition and removal of both PBS 1X and media were gently done using a multichannel pipette. Each experimental medium was freshly prepared the day of the treatment. The experiments were carried out according to the manufacturers’ instructions. After 24 hours treatment, media were collected and a dilution of 1:25 (in PBS1X) was done in order to fit the linear range of the assay indicated by the commercial kit. The diluted samples were analysed the same day or frozen at -80°C and processed afterwards. The assay was done at room temperature with media and reagent RT equilibrated to avoid uneven signals. Luminescence was recorded using Infinite M200 plate reader (Tecan). Protein content was evaluated using the BCA protein assay (Pierce). The results were presented as nmoles/minutes/grams of protein.

#### Stable isotope labelling, metabolite extraction and quantification

Cells were grown at standard culturing conditions as described in cell culture methods; the stable-isotope labelling was obtained by culturing cells with media containing either fully labelled [U-^13^C] glucose, [U-^13^C] glutamine or [U-^13^C] pyruvate for 24 h. Before metabolite extraction, plates were taken to a cold room and 500μL of medium from each plate was frozen for later analysis. The remaining media was removed, and the plates placed in Fan ice/water bath before washing twice with ice-cold PBS.

For metabolites extraction, 800μL ice-cold methanol containing an internal standard of 1mM scyllo-inositol was added to each plate and cells were detached by scraping. This mixture was transferred to a 15ml tube, and the plate washed with 800μL of methanol:H_2_O (1:1 vol/vol) that was moved to the tube, to which 400μL of ice-cold chloroform was added. The tubes were placed in a water bath sonicator in a cold room for 1 h, with 3×8 minute pulses of sonication and centrifuged for 10 minutes at 16,000rpm at a temperature of 0°C. The supernatant was extracted and dried in a vacuum concentrator. The cell pellet was then re-extracted with 300μL of methanol:H_2_O (2:1 vol/vol), this was sonicated, spun and the supernatant added to the previous supernatant tube and dried again in a vacuum concentrator. The remaining cell pellet was used for estimating dry weight and measuring total protein. For phase partitioning, the dried supernatant was resuspended in 350μL of chloroform:methanol:H_2_O (1:3:3 vol/vol) and spun for 5 minutes at 0°C and 16,000rpm. The extract is then in a biphasic partition, with the upper phase containing the polar metabolites and the lower phase containing lipidic metabolites. The polar phase fractions were then transferred to GC-MS vial inserts and vacuum-dried. Separate vial inserts had 10μL of the saved cell culture medium added, with 1mM scyllo-inositol, which were also dried in a vacuum concentrator. Each vial insert had 30μL of methanol added, containing 1μL of 5mM nor-leucine as another internal standard, followed by 30μL of methanol without nor-leucine, with the vials being dried in a vacuum concentrator after each addition. Before running samples on the mass spectrometer, polar fractions were derivatised, 20μL methoxyamine (30mg/mL in pyridine) was added to each insert and this was vortexed briefly and incubated at room temperature overnight, Silylation was then done by adding 20μL of BSTFA + TCMS reagent to each insert and incubating for 1 h at room temperature.

For metabolites analysis, an Agilent 7890A GC with a 5975C triple axis detector MSD (Agilent Technologies, Santa Clara, CA) was used. Metabolites were separated on an Agilent J&W 122-5532G DB-5ms capillary column (30m x 0.25mm, 0.25um film thickness), in splitless mode. The injector and transfer line temperatures were 270 and 280°C, respectively. The flow rate of helium carrier gas was 0.7 mL/min. The oven temperature was programmed to hold at 70°C for 2 min, increased to 295°C at a 12.5°C/min ramp rate, increased from 295°C to 320°C at a 25°C/min ramp rate, and held at 320°C for 3 minutes. The mass spectrometer was operated in scan mode, after a 6 minute solvent delay with a range of 50−565 mass/charge (m/z) and a scan-rate of 2.8 scans per second. Metabolites were identified by matching retention times and fragmentation patterns to commercially available standards. Metabolite peaks were integrated at each isotopologue m/z using MassHunter Workstation software (Agilent Technologies). Peak areas were quantified based on peak areas of known standards using nor-leucine as an internal standard, and then metabolite levels were normalised to protein content. Mass isotopologues were stripped of the contribution from natural abundance, based on the chemical formula of derivatized fragments quantified. Percent enrichment for an isotopologue was calculated by dividing the corrected intensity by the sum of corrected intensities of all isotopologues for that metabolite. Significance of metabolite enrichment between different samples was calculated with one-way ANOVA.

#### Quantification and statistical analysis

Experimental design and statistical analysis are described in the Figure legends.

## Results

### 1. Discovery of large gene cluster switches, defining breast tumour mitochondrial and metabolic subtypes

In order to identify co-regulated gene clusters in subgroups of breast cancer samples we used our previously established massive correlating biclustering method, MCbiclust^30^. The method is designed to find subsets of samples in which large genesets, such as sets of metabolic and nuclear encoded mitochondrial genes^31,32^ are co-regulated, as demonstrated by the correlation of individual gene expression levels across the selected samples. Technically, an iterative stochastic greedy search method selects the group of samples in which the chosen gene set reaches the maximum absolute correlation score. The samples and gene sets that have the highest score define a bicluster.

As an additional feature, MCbiclust using the absolute correlation values for sample selection allows the finding of two groups of genes that have maximal positive correlations within the groups, but negative correlation between each other. Thus, two genesets with mutually exclusive expression are obtained in each bicluster, defining two groups of samples (see^31^ and Figure 1A). Importantly, after defining the bicluster, quantitative correlation is extended to the (i) whole sample set using the similarity of each sample to the bicluster sample set, ranking them according to the correlation of their bicluster gene set expression with that of the bicluster samples, and (ii) to the whole transcriptome, using the correlation of each gene transcript with the average expression of a set of representative bicluster genes (correlation vector values, CV). Thus samples and gene sets can be analysed independently (for the schematics of the method see Figure 1A).

We applied the method to three independent breast cancer transcriptome datasets, METABRIC^33^, TCGA^34^ and Oslo2^35^ and obtained equivalent results. MCbiclust identified three independent biclusters in the METABRIC dataset (termed mitochondrial biclusters MB1, MB2 and MB3, Figure S1A-C), indicating three different large genesets that are strongly co-regulated (including both + or - correlation) in specific subsets of the METABRIC samples. Accordingly, each sample in the bicluster belongs to either of the two positions of a toggle switch, splitting up samples in two groups (termed upper fork, UF and lower fork, LF, see below) driven by their unique gene sets, anti-correlated between the sample groups. To visualise the switches, for each bicluster we plotted the ranking of samples (according to their correlation strength, i.e. the average CV of their bicluster genes) against the differences in the correlation in their gene sets, captured by the first principal component (PC1) of each sample (Figure 1B). The distribution pattern of the samples showed the split nature of the biclusters, with bifurcating PC1 values in highly ranked samples. The same toggle switches were confirmed in the TCGA and Oslo cohorts (Figure S1D,E).

Next, we compared the sample composition across the biclusters and explored gene sets which dominate the clusters. While most forks of the three biclusters were composed of unique samples (Figure 1C), MB1_LF and MB2_LF showed a near complete overlap, and were therefore aggregated into a common cluster (MB12_LF). To define the position of the samples in all of the discovered biclusters, we calculated their ‘forkscale’, indicating how close they are to the tip of the upper (+1) or lower (-1) forks and compared these values between the biclusters (Figure 1D, E). Remarkably, comparison between the MB1 and MB2 bicluster switches reveals a tripolar distribution where samples are merged in a common lower fork (MB12_LF) and are split between two distinct upper forks, MB1_UF and MB2_UF. This indicates the presence of a multi-state switch. Comparison of MB3 with either MB1 or MB2 did not show any significant overlap however, indicating independently regulated gene cohorts in the MB3 bicluster.

To define the gene sets dominating the biclusters we have ranked the whole transcriptome based on the CV of each gene, indicating its correlation with the switch pattern. Figure 1F shows the correlation map of the MB1 bicluster. Genes with the highest and lowest CV values were considered the dominant, most strongly regulated genes in the UF and LF switch positions, respectively. Expression levels of these genes are anticorrelated between the UF and LF samples (see Figures 1A and S1F). Importantly, genes with the highest CV values (in both UF and LF switch directions) include genes additional to the bicluster, indicating that they are part of larger scale gene regulation patterns (Table S1). The patterns and their relationship with the bicluster genes are explored in the next section.

Notably, nuclear encoded mitochondrial and metabolic genes were markedly over-represented in the UF of the MB1 and MB2 bicluster patterns but were excluded from the MB12_LF as shown by the metabolic and mitochondrial gene annotation in Figure 1F. The relative enrichment of nuclear encoded mitochondrial and metabolic genes are shown in Figure 1G.

Altogether, MCbiclust identified large scale gene expression switches defining breast cancer subtypes, which include the highly controlled co-regulation of a large group of metabolic and mitochondrial genes. The gene sets identified (see Table S1), showed switch-like behaviour, schematically represented in Figure 1H.

### 2. Defining metabolic patterns of mitochondrial breast cancer subtypes and their association with cell state

Using the gene and sample sets generated by MCbiclust, we created a pipeline to generate a series of predictions, covering tumour metabolism, oncogenic signalling and clinical outcome. In order to understand the transcriptional metabolic signatures (TMS) of the identified switches, we mapped all metabolic genes in all three switches on the global KEGG metabolic pathway map (Figure S2A). These maps indicated that the TMSs comprise all major metabolic modules, including carbohydrate, lipid, nucleotide, amino acid, glycan, cofactor and energy metabolism. Importantly, in each switch, the TMSs of the UF and LF sample sets were associated with either mutually exclusive pathways (e.g. lipid synthesis versus degradation), often with enzyme isotype switches or alternative usage of substrates (e.g. glycolytic versus glutaminolytic energy production).

To perform a more detailed *in silico* analysis of the metabolic phenotypes predicted by the method, we focused on the MB12_LF vs MB1_UF/MB2_UF bi-state switch, using two approaches. First, we compared the CVs of each gene of individual metabolic pathways, highlighting differences in their association with different switch positions. As shown in Figure 2A, in central carbon metabolism, differential correlation (i.e. difference in CVs in the upper and lower switch positions) of glycolytic and glutaminolytic enzyme transcripts demonstrated the transcriptional basis of metabolic phenotype switches, as well as the difference between MB1 and MB2 biclusters (further pathways are shown in Figure S2B).

**Figure 2.**
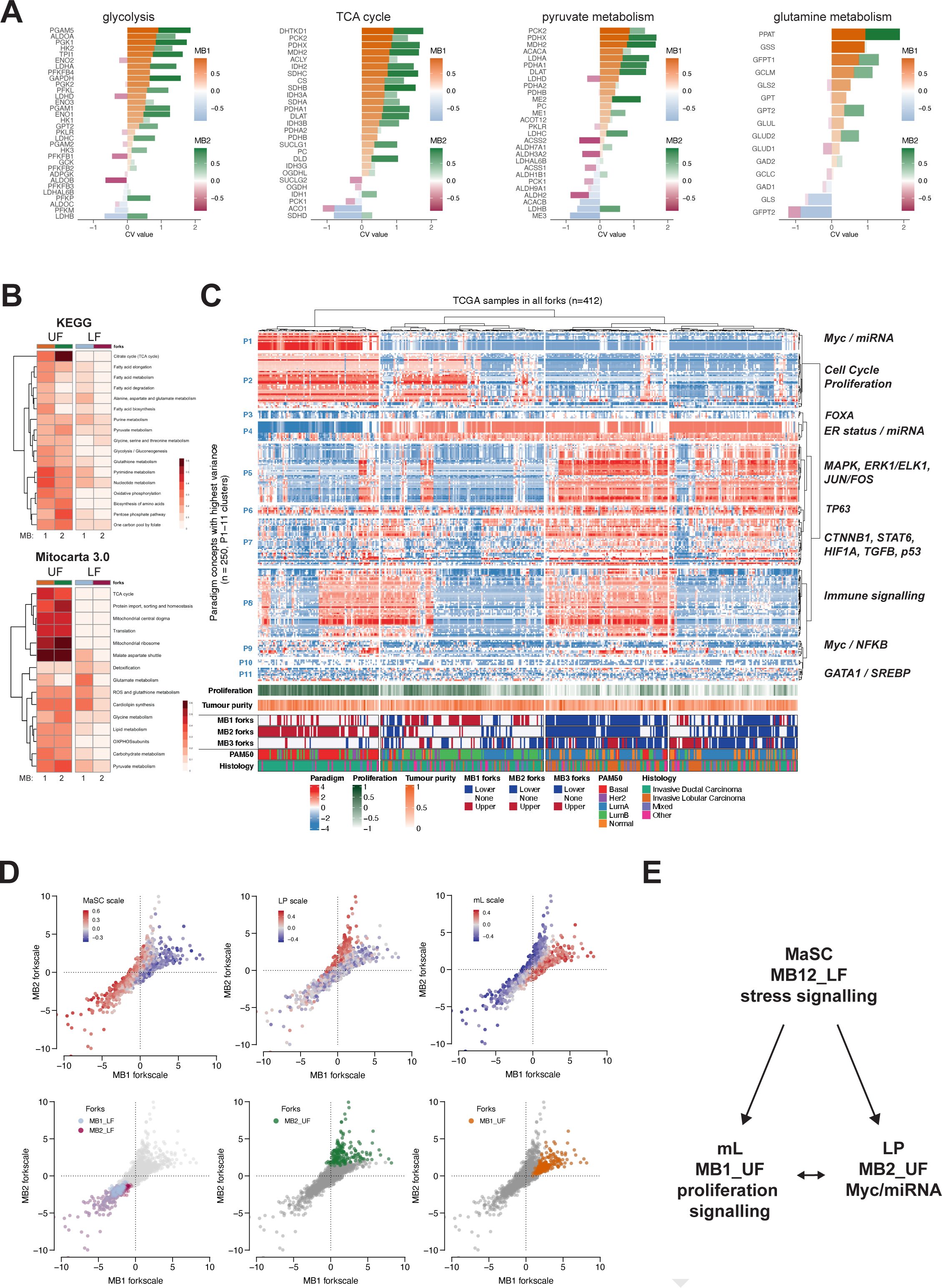
Decoding the transcriptome switch from individual genes to large scale cell state patterns. **A, B.** Metabolic phenotype predictions in the MB_1 and MB_2 bicluster switches. CV values (association with bicluster, see Figure 1 and Methods) of individual genes in the METABRIC breast cancer dataset analysis shown for pathways: glucose, glutamine, pyruvate metabolism and the TCA cycle in **A**. Values are ranked by the MB_1 CV value. The figure allows the identification of high CV genes and where differences are large between the switches (biclusters). The representation of pathways according to the relative number of genes affected is shown in **B** as a heatmap (top: based on top metabolic KEGG pathways; bottom: top MitoCarta3.0 ontology pathways, see Table S5). **C.** Heatmap of distribution of the top 250 most variable PARADIGM concepts across the bicluster switches in the TCGA BRCA dataset. The term values were obtained from ^37^ pan cancer analysis and hierarchically clustered. Annotations show proliferation, tumour purity, switch identity (UF and LF of MB_1, MB_2 or MB_3), Pam50 and histology classifications. Transcriptomic signatures based on PARADIGM concepts were assigned as shown at the right. **D, E.** Correlation of known mammary gland cell states and the transcriptomic switches in the METABRIC dataset. 2D distribution plots of samples by two biclusters is shown in **D**, overlaid by either cell of origin marker enrichment values (obtained by gene set variation analysis, GSVA - see Methods; mL: mature luminal, LP: luminal progenitor, MaSC: mammary stem cell, genesets from ^38^)

Second, to identify individual genes defining the three states, we used the CVs of the whole genome in the MB1 and MB2 bicluster switches, and identified groups of genes distinguishing the forks (CVs differ by at least 0.4 between the switches and the absolute value of the CV is higher than 0.6), as well as those common to both biclusters (Table S2). The resulting gene sets were then filtered for metabolic and mitochondrial genes used for pathway analysis (see Figure 1F, G), of which the key findings are shown in Figure 2B. Importantly, using both KEGG pathways and the MitoCarta3.0 gene set and ontology^36^, both the MB1_UF and MB2_UF switch positions showed significant activation of mitochondrial biogenesis pathways (KEGG: oxidative phosphorylation, TCA cycle; Mitocarta: mitochondrial central dogma: including mtDNA replication, transcription and translation on the mitochondrial ribosomes), as well as increased OXPHOS components (Figure S2B, Tables S3 and S4).

These results indicate that the metabolic switch between the MB12_LF and both the MB1_UF and MB2_UF positions encompass similarly increased mitochondrial biogenesis and predict higher electron flux through the TCA cycle and OXPHOS compared to MB12_LF. Whilst most of the intermediary metabolic pathways, with associated nucleotide, lipid and amino acid metabolism were more strongly represented in MB1_UF and MB2_UF, both cellular (KEGG) and mitochondrial (Mitocarta) glutamate metabolism had stronger association with MB12_LF, representing an almost uniquely associated pathway in this switch. This indicates an altered substrate preference. We noted that whilst the MB1_UF and MB2_UF positions both showed higher mitochondrial biogenesis, the pattern of correlation of individual metabolic genes differed in some key points between these switch positions. Nucleotide, fatty acid and glutamate metabolism were strongly regulated in MB1_UF, whilst there was a switch to pyruvate metabolism and TCA cycle with enhanced pentose phosphate shunt in MB2_UF. This reflects the activation of distinct metabolic switch mechanisms, presumably due to differences in cell states and their supporting metabolic configuration, driven by cell state specific signalling.

Next we asked whether the observed metabolic switch positions with divergent mitochondrial biogenesis follow the activation of specific signalling pathways. To infer signalling pathway activity in individual samples, we analysed the distribution of the results of a previously implemented PARADIGM analysis^39^ on a pan-cancer dataset^37^ among different switches. Figure 2C shows the clustering of the most divergent PARADIGM pathway entities and their association with different biclusters and forks. Overall, the switch between lower and upper fork positions is dominated by turning on a cell proliferation cluster (MYB, FOXM1, E2F family, *P2*, Figure 2C) in the upper forks (MB1_UF, MB2_UF) and by a stress and hypoxia epig(MAPK, p53, HIF1, *P5-7*) and stemness (p63) signalling cluster in the lower forks (MB12_LF). Interestingly, the MB1_UF and MB2_UF samples clustered separately, likely due to activation of Myc in MB2_UF together with a miRNA pattern (*P1*). While both MB1_UF and MB12_LF switch positions showed strong estrogen receptor signalling (*P4*), MB2_UF contained only ER negative samples (for full gene list see Table S6, for statistical analysis see Figure S3). All bicluster switches contained only a fraction of samples with high immune signature, indicating that differences between the forks is not determined by the immune system.

Taken together with the metabolic gene expression data, our model predicts higher mitochondrial content in MB1_UF and MB2_UF as compared to MB12_LF, which can be thus driven by the cell cycle and proliferation gene network activated in the upper switch positions (see Figure 2C). The differences between MB1_UF and MB2_UF can be assigned to Myc activation. Conversely, MB12_LF, showed no transcriptional (co-)activation of nuclear encoded mitochondrial genes, but rather demonstrated a clear switch in substrate preference to glutamate from glucose derived pyruvate, a metabolic phenotype associated with stress signalling.

Finally, considering that metabolic switches have been associated with cell states and cell fate decisions such as stemness and differentiation^40–42^, we assessed whether the identified metabolic and signalling switches among groups of breast cancer samples relate to their cell of origin, i.e. their state of stemness or differentiation in a specific direction. Although the so-called intrinsic (PAM50) breast cancer subtypes have been recently associated with specific states of mammary epithelial cell differentiation^43^, we have found only a loose association between switch states and intrinsic subtypes (Figure S4A). Therefore, we used an unbiased approach by calculating GSVA scores (gene set variance analysis^44^) for gene expression patterns derived from mammary stem cells (MaSC), luminal progenitors (LP) and mature luminal cells (mL)^38^, and overlaying the values of each cell-of-origin pattern for each sample on the fork distribution plots (Figures 2D, S4B). Strikingly, there was an apparent association between the three cell states and the MB12_LF / MB1_UF / MB2_UF multi-state switch (Figure 2D). Average scores for all three cell states were significantly higher than for any of the intrinsic (PAM50) subtypes (Figure S4C). Stemness (MaSC) was largely accompanied by MB12_LF features, such as low mitochondrial abundance and glutamate substrate preference, whilst switching to either mL or LP cell state was followed by increased mitochondrial biogenesis driven by proliferation signalling in absence (MB1_UF: mL state) or presence (MB2_UF: LP state) of Myc/miRNA signals (Figure 2E).

### 3. Mitochondrial and metabolic gene expression profiles predict functional metabolic switches

For functional verification of the predicted metabolic switches, we aimed to detect the identified gene expression switches in cellular and *in vivo* breast tumour models. Using either our custom or GSVA scoring methods on breast cancer cell line transcriptome datasets^45–47^, we identified cell lines belonging to each position of the three bicluster switches, demonstrating that cellular models represent appropriately the full spectrum of transcriptional signatures identified in human samples (Figure 3A, Table S7). On the other hand, in a collection of patient-derived breast cancer xenograft (PDX) models^48^, the MB1-2_LF position was not represented among the successfully implanted PDXs, although all the biclusters and forks were discoverable in the original tumours (Figure 3B). We therefore used cell line models to explore two hypotheses based on the predictions from computational analysis: cells with MB1_UF phenotype (i) boast higher mitochondrial content and activity, and (ii) have altered substrate preferences as compared with the MB1_LF phenotype. We used MCF7 and T47D cell lines to represent the MB1_UF switch and MDA-MB-436, Hs578t and HCC1134 cell lines for modelling the MB1_LF phenotype.

**Figure 3.**
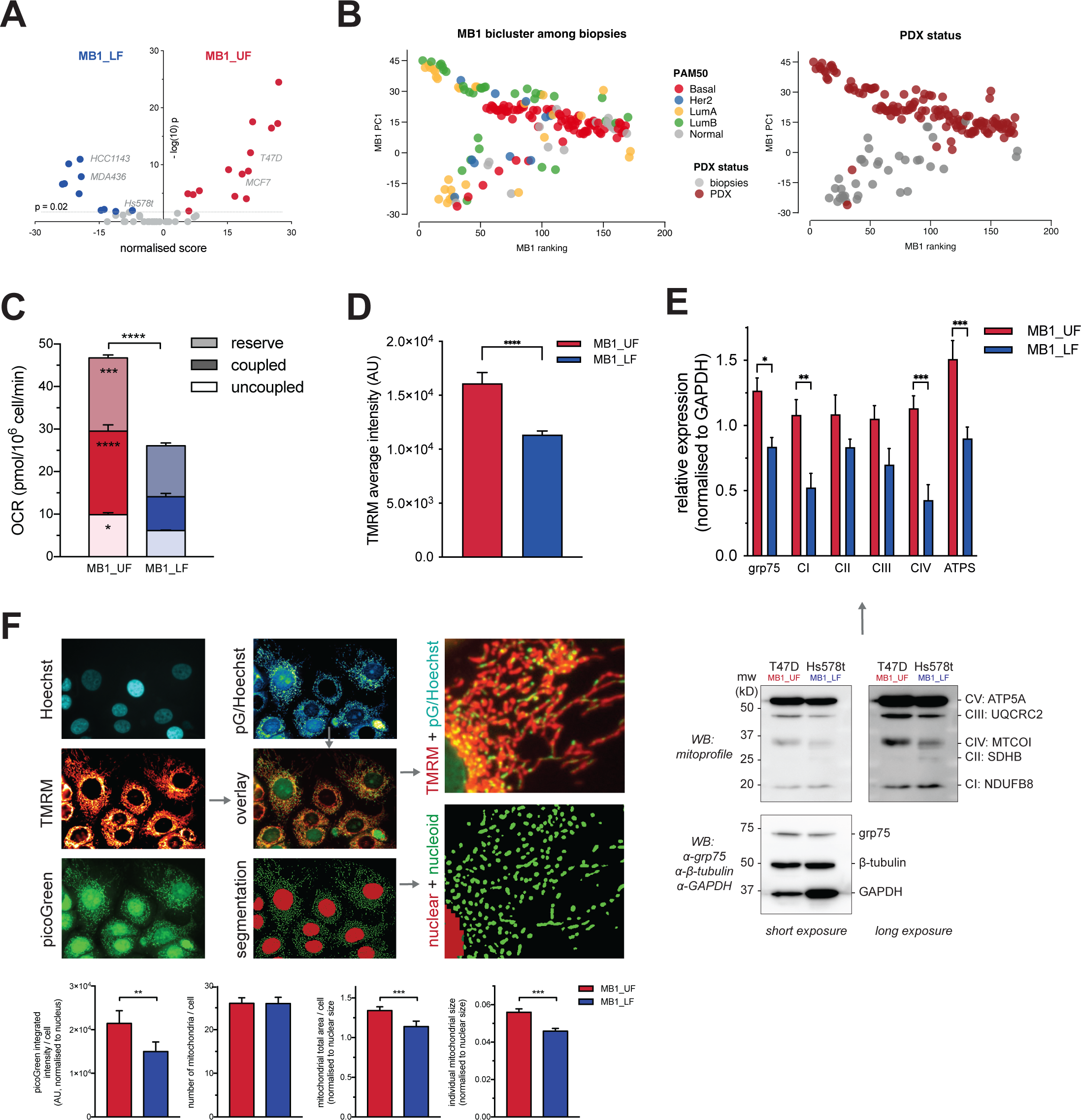
Mitochondrial function: MB1_UF cells show active functional biogenesis as compared to the MB1_LF phenotype. **A.** Cell line scoring to reveal cell lines belonging to the observed switches. Volcano plot of the scores (see Methods, Table S7) and -log_10_ of the p-values in the MB1_UF / MB1_LF switch, with the cell lines chosen for experimental work highlighted in red. **B.** 2D distribution (PC1 vs ranking index, shown for the MB1 bicluster switch) of biopsies obtained for generating PDX samples from the study ^48^, overlaid in left panel: with Pam50 categories to highlight the similarity to the MB1 switch (see Figure 1B) or in right panel: with growth success whether the sample has grown from the biopsy to PDX status. **C.** Oroboros evaluation of ∼1M cells of the cell lines grouped by their switch state (MB1_UF / MB1_LF). Reserve, ATP synthesis coupled and uncoupled respirations were quantified using FCCP and oligomycin treatments (See Methods). 2-way ANOVA with Sidak’s multiple comparisons test. **D.** Average cellular TMRM intensity grouped by switch state (MB1_UF / MB1_LF), data obtained from high content microscopy from > 1K cells belonging to either switch state. Unpaired Student’s t-test. **E.** Relative expression of respiratory chain subunits quantified from western blot analysis. Results are grouped by switch state (MB1_UF / MB1_LF). 2-way ANOVA with Sidak’s multiple comparisons test. **F.** High content imaging analysis of mitochondrial biogenesis, structure and function in MB1_UF and MB1_LF cells. Cells were loaded with picoGreen, Hoechst and TMRM and imaged on an ImageXpress Micro XLS widefield high content screening system, as described in Methods. Hoechst normalised picoGreen signal, for amplification of the mitochondrial picoGreen signal, was overlaid and segmented for quantitative analysis on the TMRM signal, denoting thus nucleoid (green) and mitochondrial (red) segments. Below the images, quantifications in cell lines grouped by their switch state (MB1_UF / MB1_LF) of the picoGreen integrated intensity, readout for mtDNA content (left panel); of mitochondrial number, total volume and individual mitochondrial volume, to assess the state of the mitochondrial network (three panels to the right, respectively). Unpaired Student’s t-tests. (ns: P > 0.05, *: P ≤ 0.05, **: P ≤ 0.01,***: P ≤ 0.001, ****: P ≤ 0.0001)

In order to determine the differences in mitochondrial content and activity, we measured oxidative mitochondrial function and used high-resolution, high-content imaging on large cell populations to quantify overall mtDNA content and morphology in the MB1_UF and MB1_LF fork cell lines (Figure 3C-F). MB1_UF cells had higher oxygen consumption rate and mitochondrial membrane potential, indicating higher redox potential to feed electrons in the respiratory chain (Figure 3C, D). Higher mitochondrial abundance in MB1_UF cells was confirmed by western blot analysis of respiratory chain components (Figure 3E). High-resolution image analysis of mitochondria across cell populations (Figure 3F) showed that while the number of mitochondria per cell was equal in both forks, MB1_UF cells had higher mtDNA content, and greater total and individual mitochondrial area, indicating larger mitochondrial size and overall content. These experiments confirmed a switch in mitochondrial abundance and function as predicted by gene expression profiling using MCbiclust.

Next, in order to detect the metabolic switch with altered substrate preferences associated with the different mitochondrial switch positions, we performed a steady state metabolic flux analysis of heavy carbon labelled key cellular fuels using uniformly labelled ^13^C-glucose, ^13^C-glutamine and ^13^C-pyruvate. MB1_UF and MB1_LF cells were incubated with each substrate in separate experimental sets. Mass spectrometry analysis of carbon-labelled compounds was used to quantify isotopologues of metabolites in the main catabolic pathways of the three substrates. In addition, to verify substrate dependencies, we introduced conditions with combinations of reduced glutamine and pyruvate (glutamine[glutamine] H/L = 1mM/0.1mM; [pyruvate] +/- = 1mM/0 mM, Figure 4A).

Using these methods, the fate of glucose via aerobic glycolysis was assessed by quantifying glucose uptake, lactate production, and the relative incorporation of glucose derived heavy carbons into lactate (m+3 isotopologue). We found no significant differences between the MB1_UF and MB1_LF switches in all the conditions tested (Figure 4B). This shows that aerobic glycolysis occurs in both MB1 switches at high rate, verified by the nearly complete glucose-derived ^13^C labelling of glycolytic intermediates (Figure S5A). However, MB1_UF cells produced pyruvate (m+3) from glucose at significantly higher levels than MB1_LF (Figure 4C), suggesting that glucose flux through glycolysis was higher in the MB1_UF switch, presumably delivering pyruvate to be oxidised in the TCA cycle.

The TCA cycle can be fed either by glucose derived pyruvate or glutamine derived glutamate at any ratio (Figure 4D). Thus, to assess substrate preferences, we quantified the relative carbon contribution of glucose, glutamine and externally added pyruvate carbons to the TCA cycle. The results showed significantly higher glucose-derived pyruvate incorporation into citrate, fumarate and malate in the MB1_UF cells, while more glutamine derived carbons were incorporated into TCA cycle metabolites in MB1_LF cells (Figures 4E, S5B). These results indicated a clear switch between glucose-derived pyruvate versus glutamine feeding into the TCA cycle. Pyruvate entered the TCA cycle both via the PDH and PC routes in MB1_UF cells (Figure 4F). In contrast, MB1_LF cells used glutamine in both the oxidative and reductive direction at higher levels than in MB1_UF cells, and additionally showed increased glutamine uptake and subsequent glutamate secretion (Figure 4G). In summary, both low (MB1_LF) and high (MB1_UF) mitochondrial activity switches in cultured tumour cells are accompanied by aerobic glycolysis, but the switches differ in their substrate preference to fuel their TCA cycle activity. The MB1_UF switch promotes the use of glucose as main carbon source, while in MB1_LF glutaminolysis dominates.

To functionally verify the switch to glutamine preference in the MB1_LF phenotype, we applied glutamine restriction (see conditions in Figure 4A). The lack of glutamine resulted in reduced bioenergetic competence demonstrated by reduced mitochondrial oxygen consumption rate and total cellular ATP levels limited to MB1_LF cells (Figure 4H). Strikingly, glutamine restriction partly reverted the MB1_LF metabolic switch in the MB_UF direction by promoting glucose derived pyruvate entry in the TCA cycle via the PC route (see Figure 4F). Interestingly, externally added pyruvate was incorporated at only low levels into the TCA cycle (Figure S5D) but promoted glucose derived pyruvate utilisation by the PC route in MB1_LF cells, shifting the metabolic phenotype towards the MB1_UF switch (Figure 4F).

**Figure 4.**
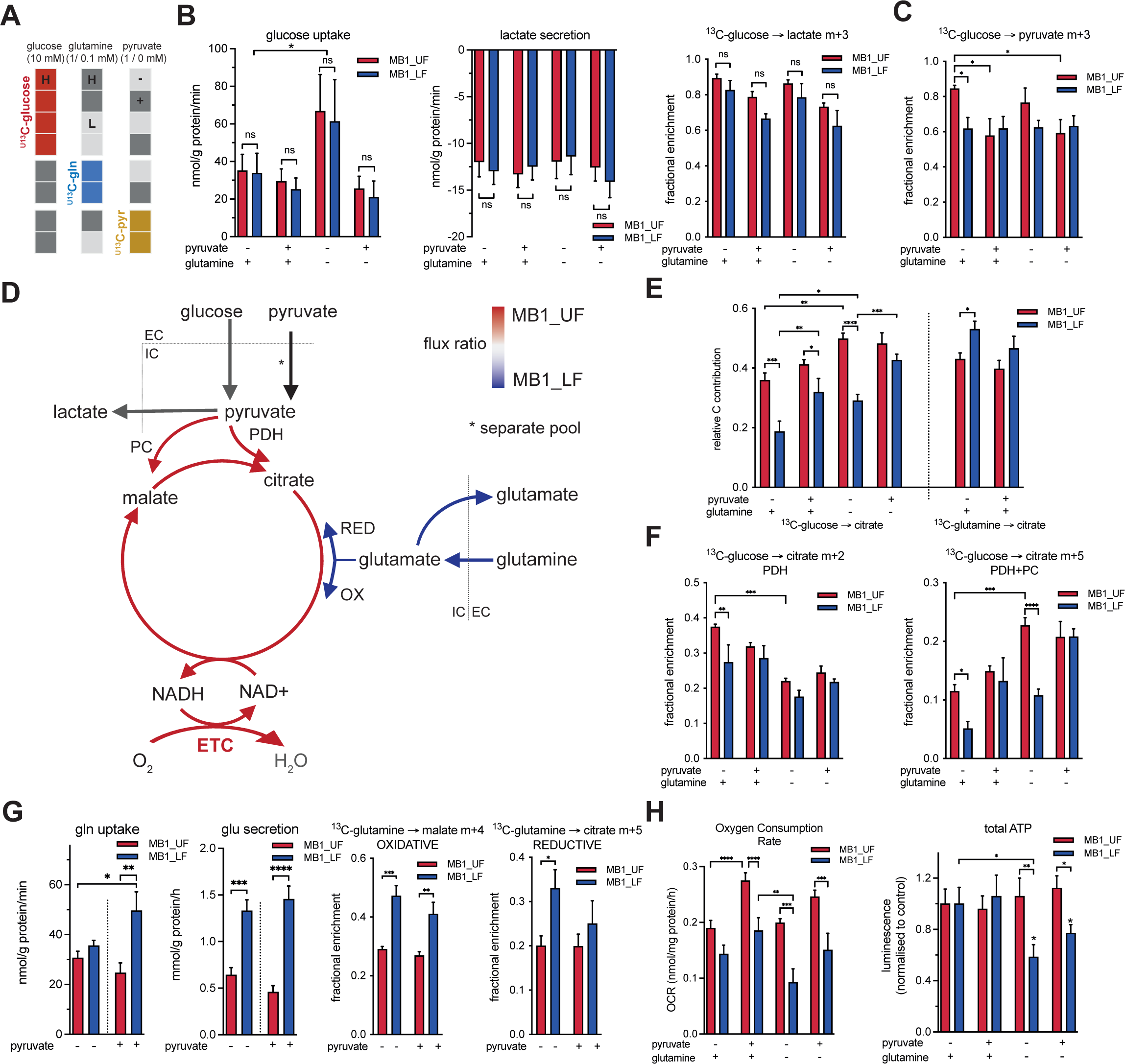
Metabolic wiring of the switch positions probed by 13C-labelled substrates. **A.** Nutrient supply and ^13^C-labelling conditions used throughout the experiments presented in this figure. High and low glutamine (H/L = 1mM/0.1mM) levels were combined with the presence and absence of pyruvate (+/- = 1mM/0 mM), glucose was kept at 10mM. **B.** Aerobic glycolysis was tested by measuring glucose uptake (first panel) and lactate secretion (second panel) from the mass spectrometry data obtained from cell culture media, as well as the fractional enrichment of m+3 labelled lactate intracellularly (third panel), see Methods. Cells were grouped by switch state (MB1_UF / MB1_LF). 2-way ANOVA with Sidak’s multiple comparisons test. **C.** Fractional enrichment of the m+3 pyruvate isotopomer, following ^U13^C-glucose labelling for 24h, in cells grouped by switch state (MB1_UF / MB1_LF). 2-way ANOVA with Sidak’s multiple comparison tests. **D.** Overall model of the MB1_UF/MB1_LF metabolic switch in central carbon metabolism. Glucose derived pyruvate is preferentially used in the TCA cycle either by the PDH or PC route in the MB1_UF cells/position, driving high respiration. Glutamine is taken up more in MB1_LF cells, and they use it preferentially in the TCA cycle, and their respiration relies on glutamine derived glutamate, but glutamate is also re-secreted into the extracellular space in high amount in these cells. Extracellular pyruvate does not contribute significantly to the TCA cycle, preferentially used for lactate production and thus setting the cytosolic redox ratio. **E.** Relative carbon contribution to citrate following 24h labelling with ^U13^C-glucose and ^U13^C-glutamine, in cells grouped by switch state (MB1_UF / MB1_LF) The values were calculated as described in Methods. 2-way ANOVA with Sidak’s multiple comparison tests. **F.** The fate of glucose derived carbons in the TCA cycle. Fractional enrichment of the m+2 (PDH route, left panel) and m+5 (PDH+PC route, right panel) isotopologues in citrate following 24h labelling with ^U13^C-glucose, in cells grouped by switch state (MB1_UF / MB1_LF). 2-way ANOVA with uncorrected Fisher’s LSD multiple comparison tests. **G.** The fate of glutamine in cells grouped by switch state (MB1_UF / MB1_LF). Glutamine uptake and glutamate secretion rates were measured by specific biochemical assays in the culture media prior and following 24h incubation with cells (left two panels). Fractional enrichment of malate m+4 (oxidative route) and citrate m+5 (reductive carboxylation) isotopologues following 24h labelling with ^U13^C- glutamine (right two panels). 2-way ANOVA with uncorrected Fisher’s LSD multiple comparison tests. **H.** Substrate dependence of mitochondrial respiration and cellular ATP production in cells grouped by switch state (MB1_UF / MB1_LF). Left panel shows basal oxygen consumption rate as measured by Seahorse XFe96 analyser (see Methods). Right panel shows total cellular ATP content measured by a luciferase assay following lysis of cells kept in the indicated media for 24h. 2-way ANOVA with uncorrected Fisher’s LSD multiple comparison tests. (ns: P > 0.05, *: P ≤ 0.05, **: P ≤ 0.01,***: P ≤ 0.001, ****: P ≤ 0.0001)

Thus we have discovered a metabolic switch defining MB1_UF tumour cells where a mature luminal (mL) cell-of-origin phenotype with oestrogen receptor and cell cycle signalling is associated with an adaptive gene expression pattern, promoting high glycolysis together with high TCA activity and energy production, relying mostly on glucose. In the other position of the switch, in MB1_LF mammary stem cell like cells (MASCs), glutamine is preferred to use to fuel lower mitochondrial activity. In addition, our functional analysis also revealed that the functional metabolic switch is adaptable to environmental changes in that the positions can be shifted by altering substrate availability. A switch from glutamine to pyruvate supply in MB1_LF cells not only reverted the low pyruvate entry into the mitochondria and corresponding TCA cycle flux to levels found in MB1_UF cells (Figure 4E, F), but also increased respiration and ATP levels (Figure 4H).

Our prediction assumes that transcriptomic profile switches can determine metabolic routes via modulating the protein expression levels of metabolic pathway components. In order to test this assumption, we performed western blot analysis of enzymes from key pathways by which glucose, glutamine and pyruvate are fed into the TCA cycle (Figures 5A, S6). Expression levels of 29 metabolic enzymes with diverse localisation clustered according to switch positions (UF and LF) across the cell lines (Figure 5B). Higher mitochondrial abundance in the UF was confirmed by increased grp75 expression compared to the cytosolic β-actin marker (Figure 5B, lower panel). Interestingly, these changes when compared with mitochondrial and cytosolic abundance, showed divergent patterns of differential expression (Figure 5B left and right heatmaps, respectively). Moreover, scaled protein levels of key enzymes matched the CV value of their transcripts (Figure 5C). Overall, expression of most metabolic enzymes followed the higher mitochondrial abundance in the MB1_UF state. Those with the largest observed difference are involved in the malate-aspartate shuttle (GOT1, GOT2) and pyruvate utilisation (MPC2, pPDH ratio, IDH2) (Figure 5D, E). These results support the metabolic phenotype described for MB1_UF cells (see Figure 4). In contrast, only a small set of enzymes was overexpressed in the LF, underpinning key characteristics of the LF, such as increased glutamine utilisation (GLS1, GFPT2). These data confirmed that transcriptomic patterns can directly predict metabolic pathway activities. Close correlation of transcript and protein levels indicated that transcriptional regulation dominates metabolic pathway activities via the regulation of their key enzymes.

**Figure 5.**
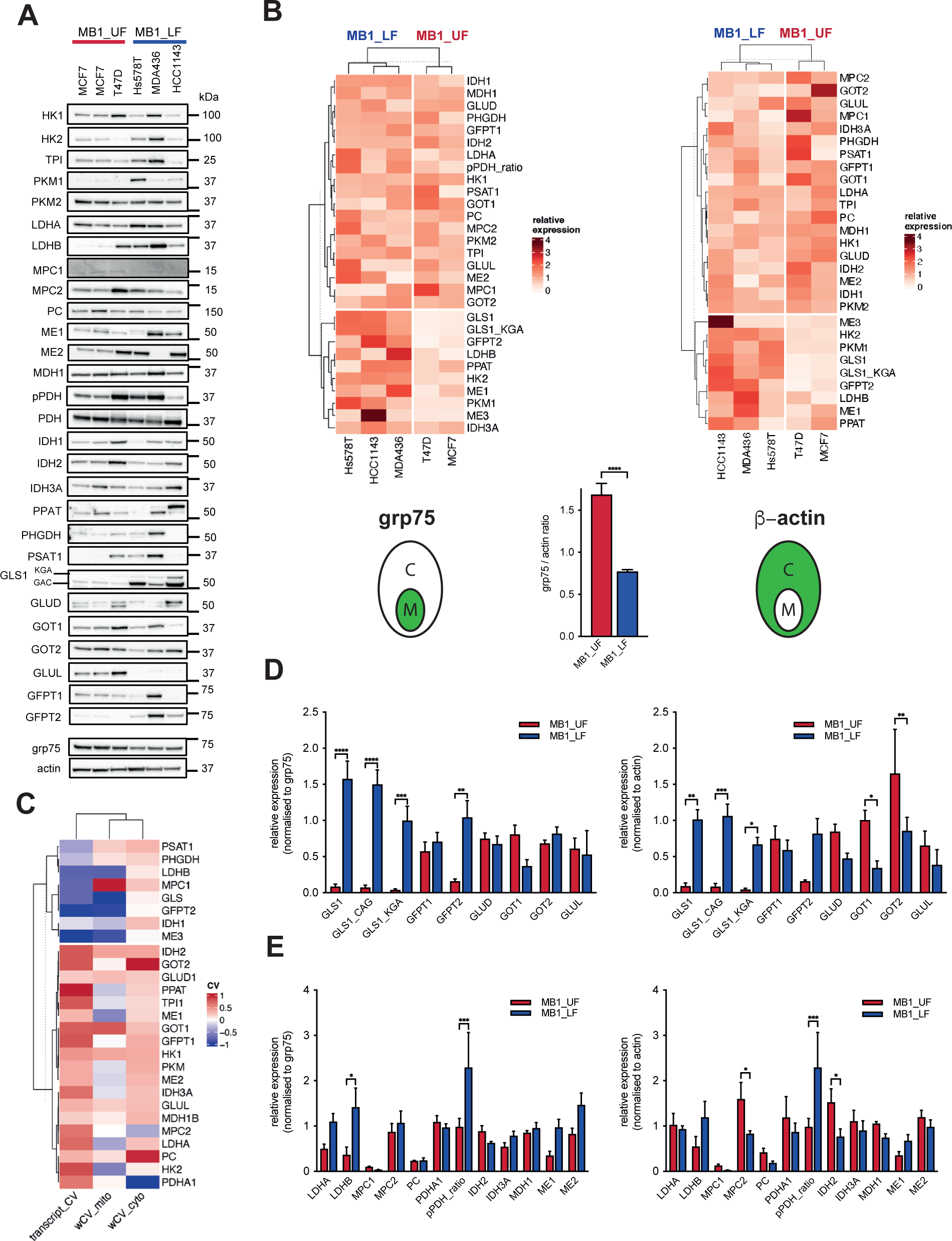
Protein expression profiles match transcriptomic switches. **A.** Western blot analysis of 29 metabolic enzymes across five cell lines representing MB1_UF and MB1_LF switch positions and cell states. Representative blots from n ≧ 3 independent experiments, used for quantification of signals, normalising density values to either grp75 or β-actin. **B, D, E.** Clustering and quantitative analysis of enzyme expression levels obtained from **A.** Values were separately normalised to either mitochondrial (left panels) or cytosolic (right panels) markers, grp75 or β-actin. The comparison of mitochondrial abundance based on the grp75/β-actin ratio is shown in the lower panel in **B.** Heatmaps are based on a pairwise, robust to outliers distance measure of the mean protein expression values. K-means clustering. **D, E.** 2-way ANOVA with uncorrected Fisher’s LSD multiple comparison tests. **C.** Heatmap and k-means clustering of (i) transcript CV values of the 29 enzymes from the METABRIC dataset (see Figs. 1, 2, positive values predicting higher expression of genes in MB1_UF, negative values corresponding to higher expression in MB1_LF), (ii-iii) the scaled (between -1 and 1) protein expression levels averaged across all cell lines, normalised to grp75 (ii) or β-actin (iii). (ns: P > 0.05, *:P ≤ 0.05, **: P ≤ 0.01,***: P ≤ 0.001, ****: P ≤ 0.0001)

### 4. Pathological and clinical predictions from metabolic gene switches

As metabolic gene switches are associated with cell states, they potentially indicate clinically relevant features such as genotypes, phenotypes and gene transcription patterns. We tested the correlation of switch positions with genetic, biological, pathological and clinical data in the TCGA and METABRIC datasets to investigate this. First, in order to understand whether a specific tumour architecture or histological origin is associated with switch positions, we used the most recent dataset with comprehensive coverage of histological features and classifications in the full TCGA BRCA sample set^49^, and tested the distribution of features among individual forks (Figures 6A, S7, Table S8). A group of features was highly variable between switch states, including enrichment for invasive lobular carcinoma (ILC) in the MB12_LF switch state, and increased signs of nuclear pleomorphism in the MB2_UF (Figure 6B). Of note, ILC, the most common “special” histological subtype of breast cancer which comprises approximately 10% of all breast cancer diagnoses in the developed world, characteristically displays low nuclear pleomorphism^50^. Overall, both the MB1_UF and MB2_UF switch positions were associated with increased proliferation (mitosis) and higher cellularity (epi_area) in contrast to MB12_LF, in which about half of tumours were classified as ILC. Additionally, MB2_UF was highly enriched in samples with necrosis, nuclear pleomorphism and immune infiltration (inflam), all of which are features of more aggressive tumours.

**Figure 6.**
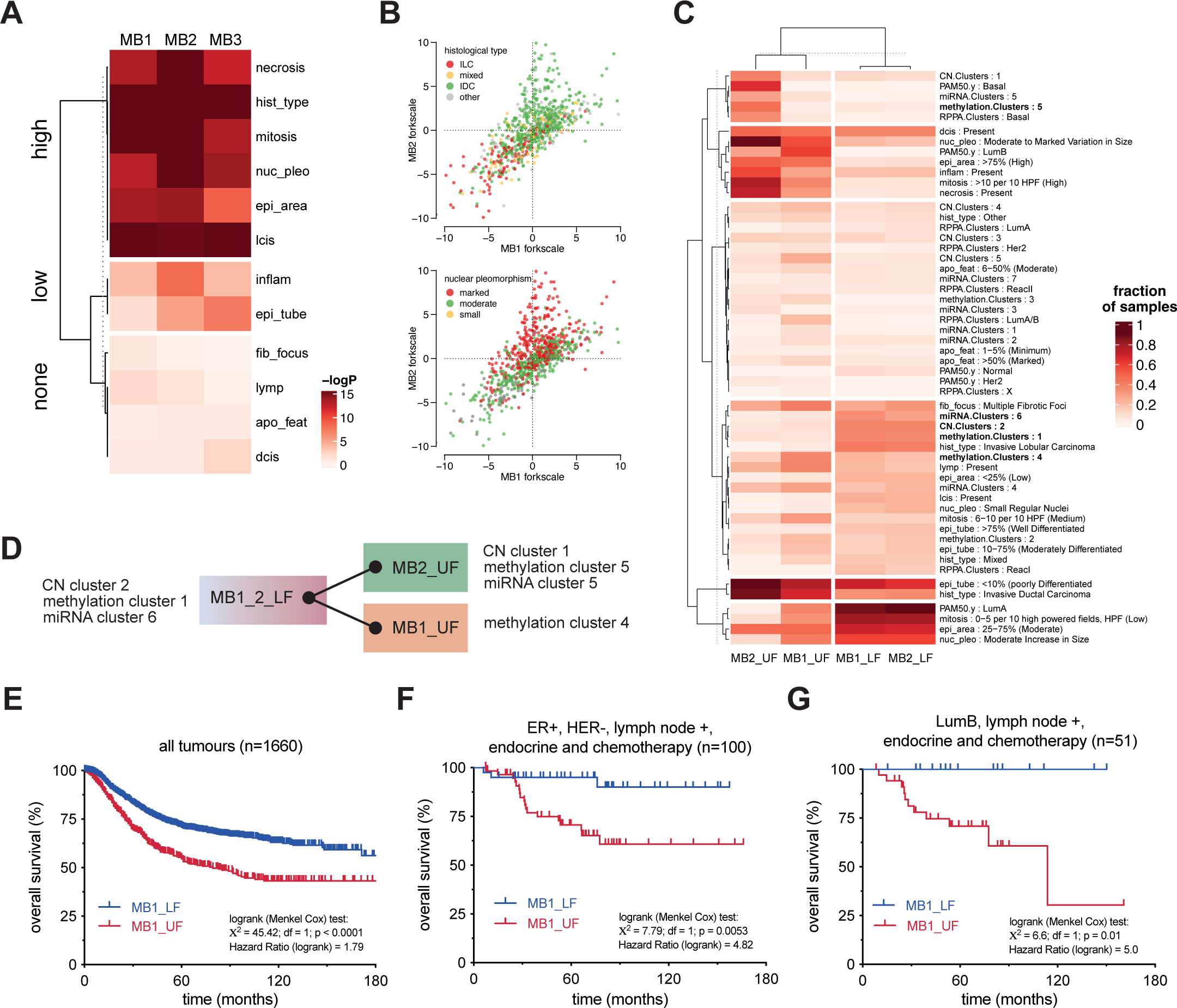
Pathological and clinical predictions from metabolic gene switches. **A.** Chi-squared distribution of discrete histological features across the upper, lower and non-fork samples of the three biclusters (MB1-MB3). Heatmap shows k-means clustering of chi-squared test p-values (-log10P), dividing the histological parameters into three groups: high, low and none association with the bicluster. **B.** Visualisation of histological feature distribution on the MB1 and MB2 switches. Classifiers of histological tumour type (upper panel) and degree of nuclear pleomorphism are overlaid onto 2D distribution plots of TCGA samples. The axes represent the scale between the upper (UF) and lower (LF) forks of each bicluster. **C.** Clustering of histological and genetic features of samples belonging to the MB1 and MB2 switches. Fractions of samples for each feature were calculated and clustered with a custom robust to outliers distance measure and K-means clustering. Genetic features representative of each switch position are highlighted in bold. **D.** Key genetic and transcriptional architecture of the MB1 and MB2 switches. **E-G.** Survival analysis of samples in the MB1 switch. Kaplan-Meier survival plots of samples with MB1_UF and MB1_LF patterns are shown. **E.** All (1660) samples compiled from the GEO database ^53^. **F.** Subset of samples from endocrine and chemotherapy treated ER positive patients (100). **G.** Subset of samples with LumB molecular subtype (51). Menkel Cox logrank test results are shown on the individual plots.

Next, in order to identify and compare further unique features of each switch position, we have assessed the fraction of samples with each individual feature spanning both histological and genetic data. As shown in Figure 6C, when analysing the MB1 and MB2 bicluster switches, the MB12_LF, MB1_UF and MB2_UF samples formed separate clusters. MB12_LF samples characteristically showed low proliferation and were enriched in tumours with the LumA intrinsic subtype and ILC histology. This is likely determined by specific CN variation, methylation and miRNA clusters (Figure 6D). Epigenetic modifications triggering metabolic switches and *vice versa*, have been recently studied^51,52^. On the other hand, while both MB1_UF and MB2_UF samples showed increased proliferation, they diverged by intrinsic subtype (LumB and Basal, respectively) and the underlying genetic and transcriptional constitution (CN variation, methylation and miRNA clusters).

Finally, we asked whether metabolic switch positions in the MB1 bicluster can predict the clinical outcome of ER positive breast cancer. In order to do this, we performed a survival analysis using an independent breast cancer dataset^53^ and applying the weighted average of a core geneset of 44 genes to determine the MB1_UF and MB1_LF switch positions. The MB1 switch includes almost exclusively ER positive tumours, divided into two groups, which partly overlap with the two luminal intrinsic subtypes, LumA and LumB and the normal-like subtype (see Figure S4A). ER positive tumours overall are sensitive to antihormonal therapy, but adjuvant chemotherapy is usually recommended against tumours progressing to lymph nodes. However, only a fraction of these patients respond to this distressing treatment, motivating large recent efforts to develop gene expression pattern based methods to stratify individual ER positive tumours into chemotherapy sensitive and resistant groups prior to adjuvant treatment^54,55^. The overall survival of patients with MB1_LF pattern expressing tumours was better than the MB1_UF group (Figure 6E). Moreover, when only tumours with the common ER positive, Her2 negative receptor subgroup from patients with axillary lymph node metastases who had been treated with both chemotherapy and endocrine therapy were considered, the 10-year overall survival of patients with the MB1_LF pattern approximated to 90% (Figure 6F). This is despite the very well-documented adverse prognosis of axillary metastases^56^, a feature that is largely independent of tumour biology. Excellent survival was also observed when the group was further filtered to include only tumours with the LumB intrinsic subtype, which is also an adverse prognostic feature^57,58^ (Figure 6G). In contrast, the survival of patients with MB1_UF tumours was dramatically poorer in these two groups.

Altogether, these results indicated that large scale mutually exclusive gene expression clusters govern the mitochondrial and metabolic status of breast tumours. These multi-state switches extend beyond the metabolic gene sets, and reveal the biochemistry, architecture and chemosensitivity of tumours.

## Discussion

The current availability of large scale genomic and transcriptomic datasets on physiological and pathological conditions escalated the need to devise computational strategies providing statistically sound predictions of how individual genes and gene groups are regulated and contribute to cellular phenotype. The Gene Expression Patterns (GEPs) underlying phenotype or cell state switches^59–61^ are complex and have not been comprehensively analysed, although methods to address the question are being developed^62^. These methods are part of pattern recognition, subgroup discovery and dimensionality reduction techniques that often apply biclustering approaches to enhance the probability of discovering patterns of gene expression which only exist in sample subsets. We have recently developed a biclustering method, MCbiclust^30,31^, which is optimised for discovery of large scale anti-correlating patterns to stratify large heterogeneous sample collections. This allows the detection of differences and switches in large-scale GEPs. We have previously shown that this approach is able to distinguish cancer cell types according to their organ (or cell type) of origin, and have discovered subgroups of mitochondrial succinate dehydrogenase (SDH) deficient tumour samples partly by their altered metabolic networks^30^.

Here we have developed a resource to apply massive correlated biclustering as part of a pipeline to discover metabolic switches and determine their signalling and transcriptional regulation, as well as their potential predictive power in tumour biology and clinical outcome. Predictions of tumour biology, energetics and metabolism were verified in experimental models and clinical relevance was demonstrated by applying the pipeline to large breast cancer transcriptome and genomic datasets. The pipeline consists of MCbiclust and a series of pathway analysis approaches used to understand metabolic adaptation in >3K heterogeneous breast cancer samples. The method identified large multi-state gene expression pattern switches in signalling and metabolic adaptation, which coincided with differentiation status, i.e the cell of origin of tumour cell types, and predicted metabolic fluxes, histological features and clinical outcome.

Currently available biclustering methods have been productive in finding functionally relevant gene networks but they do have limitations^63^. Our previous evaluation of MCbiclust^30^ showed that the algorithm scored better than five mainstream biclustering methods that were available at the time, due in particular to its ability to recognise large biclusters. Cell fate is thought to be more often regulated by widespread gene expression patterns than by activation of a small set or single genes^64–66^. This probably explains the ability of our method to discover patterns in tumour subsets that have different cells of origin. In addition, while previous methods mostly used either differential gene expression comparisons with normal samples or pairwise differential correlation^17^, MCbiclust uses a greedy search to maximise the average *absolute correlation value* of a large (∼1K) geneset. By covering a large fraction of the whole transcriptome, this allows the discovery of composite chequer pattern biclusters, which contain both positively and negatively correlated gene groups, thereby defining two gene sets that switch expression between sample subsets. The method additionally allows the discovery of alternative, multi-state switches which we have demonstrated here through our discovery that some subsets of multiple biclusters (MB12_LF) showed near complete overlap.

The relationship between transcripts of individual enzymes and metabolism is not linear^26^, and apart from enzyme transcript abundance, metabolic activity is fundamentally determined by substrate fluxes^15^. However, we have to emphasize that we do not seek to establish such a direct, linear link. Rather, we argue that long term adaptive regulation of metabolism according to cell state must be reflected by the co-regulation (correlation) of larger fractions of the transcriptome, which necessarily includes metabolism related genes. Using MCbiclust we discovered such large-scale gene expression patterns, involving a significant (4-5%) fraction of the genome. These GEPs couple cellular states via cellular signalling to cell cycle, stress responses, metabolism and bioenergetics, as shown in Figure 2C, D. We found that cellular states along the differentiation tree of mammary epithelial cells (Figure 2D, E), are represented by large GEPs which undergo binary, or multiple switching (Figure 1A-H). We argue that the mitochondrial (bioenergetic) and metabolic components of these binary GEPs carry information on the prevailing metabolic phenotype. Instead of seeking direct correlation between metabolome and transcript abundance, we assess the link in a novel way:

1. For gene group discovery, instead of mRNA abundance, we select groups of genes which show maximum correlation. This reflects strong co-expression/co-regulation among the genes in the group.
2. For the selection of the gene group members we maximised the absolute correlation value across the group, thus we considered both negative and positive correlations. This approach results in finding two groups, the most positively and the most negatively correlating ones. This means that, as shown in Figure 1A, the samples of the bicluster will have higher expression of either one or the other gene groups, as compared between each other. Accordingly, switching signifies the concerted change from ‘off to on’ (low vs high mRNA abundance) and ‘on to off’ of large (0.5-1K) groups of genes, causing the positive and negative checker pattern of correlations.
3. Since the correlation could be calculated for the whole transcriptome (Figure 1F), it was straightforward to quantify the correlation of each transcript in a metabolic pathway with the prevailing pattern (Figure 2A), giving a ranked distribution of the pathway members associating with the GEP, primarily indicating whether the correlation values are higher in one or the other sample group (upper or lower forks, see Figure 1B-E). This difference usually coincides with the difference in overall abundance of transcripts in a pathway, but the quantification is based primarily on correlation.

Thus, here we predicted relative differences in pathway activities between two groups of samples (UF and LF)in a bicluster, based on overall transcriptional activity and regulation in the groups along the specific pathway. We present experimental verification of one prediction. In this specific case, correlation analysis indicated differences in glucose and glutamine metabolism between MB1_LF and MB1_UF samples, which were confirmed experimentally (Figures 3 and 4). Previously we have also identified an MB2_UF associated metabolic phenotype, which has been experimentally verified and published^67^.

Breast cancer GEPs have been intensively studied which has led to the development of several molecular classifications; these include intrinsic subtypes^57^ as defined by the PAM50 algorithm (which is available for clinical use)^58^, and the integrated clusters which are defined using both GEPs and genomic data^33,68^. However, the further integration of the intrinsic subtypes and integrated clusters with histological data^49^ and the exact origins of other emerging subtypes are still debated^35,69^. The stratification of breast cancer provided by MCbiclust only partially overlaps with other molecular classifications such as the intrinsic subtypes (see Figure S4). It therefore provides a novel perspective on breast cancer subgroups, particularly in respect of functional features, such as metabolic and bioenergetic rewiring. This is a unique feature of MCbiclust in comparison to other molecular classifications which are all based on gene sets with much more diverse function. Importantly, other recent biclustering approaches to breast cancer datasets provide resolution such as that of PAM50 subtyping^70,71^.

Our analysis of patient survival according to the MB1 MCbiclust fork (see Figure 6) provides preliminary data to suggest that this classification may prove to be clinically very important. Although the analysis was performed in a retrospective cohort which allowed only limited matching between the characteristics of a comparatively small number of included patients, the outcome difference between the MB1_LF and MB1_UF groups is too large to have arisen by chance. The data additionally suggest that the survival prediction may be superior to that provided by intrinsic subtyping. ER positive tumours included in this analysis are sensitive to endocrine therapy as a group. The additional use of adjuvant chemotherapy has been widely recommended for patients with axillary lymph node involvement. However, only a fraction of patients benefit from this distressing treatment, motivating large recent efforts to develop gene expression pattern based methods to stratify ER positive tumours into chemotherapy sensitive and resistant groups^54,55^. The excellent outcome of patients in the MB1_LF despite other adverse prognostic features raises the possibility that patients in this group could safely avoid chemotherapy. In contrast, the poor outcome of the MB1_UF group demonstrates a need for additional or alternative approaches to treatment for these patients.

MCbiclust also provides two novel and interesting insights into tumour origin and architecture. First, by comparing previously described patterns associated with the putative cell of origin of different tumour types, a close correlation with the biclusters was found (see Figures 2D,E; S4B), illustrating the power of the method to determine cell fate associated gene expression patterns. In this analysis, the weakest correlation was found between MaSC and MB12_LF. This might not be surprising given that MaSC represents an already heterogeneous population of cells^43^, of which certain fractions might belong to other switches (in particular MB2_UF). Second, histological analysis of samples belonging to MB12_LF switch showed striking enrichment in invasive lobular carcinoma (ILC), a comparatively infrequent histological tumour subtype which is increasingly identified as an entity with molecular characteristics that are distinct from those of the common invasive ductal carcinoma^72–74^. The signalling pathways associated with the MB12_LF switch might help to explain the properties of ILC.

Intriguingly, all biclusters (including the ones arising from random genesets as starting points - see Figures 1, S1) were significantly enriched in nuclear encoded mitochondrial genes. This may reflect co-regulation of a large set of genes belonging to the same organelle and indicate the necessity of shaping mitochondrial gene expression patterns according to the actual cell state. Moreover, gene expression and metabolomic analysis have also disclosed strong co-regulation of mitochondrial and extra-mitochondrial metabolic genes. This provides a platform for understanding metabolic and bioenergetic adaptation in cancer development. Interestingly, both the MB1_UF and MB2_UF switch positions showed increased transcriptional biogenesis of mitochondria as compared to MB12_LF, where nuclear encoded mitochondrial genes were under-represented. This observation is in line with the currently emerging view of mitochondria in tumours, recognising that normal or high activity of certain mitochondrial metabolic pathways often drives aberrant tumour proliferation^75^. Our analysis focusing only on tumour samples, reveals only relative differences in mitochondrial biogenesis between the switch positions, and thus cannot discriminate between (i) upregulation of biogenesis in the MB1_UF and MB2_UF switch positions and (ii) suppression of mitochondrial biogenesis and function in the opposite (MB12_LF) position. Recent data indicating loss of mitochondrial mass and function under chronic stress conditions^76^ and reduced mitochondrial activity in stem cell states^77^ support the latter view.

In summary, we have created a methodology with the potential for discovering large, alternating gene expression patterns and have demonstrated its utility through the analysis of breast cancer cells and transcriptomic datasets. The method provides a useful resource for exploring physiological and pathological^78^ conditions where phenotypic switches play a major role.

## Acknowledgments

We thank for valuable discussions with C. Barnes, M. Duchen (University College London), K. Bianchi, C. Chelala (Barts Cancer Institute), for the support from all members of the M. Duchen, R. Rizzuto (University of Padova), MY, GS labs, for the PARADIGM data provided by R. Akbani (MD Anderson Cancer Center), and for the support from the Bioinformatics (G. Kelly) and Metabolomics (J. Macrae) STPs of The Francis Crick Institute. GS was supported by the following grants: AIRC (Associazione Italiana per la Ricerca sul Cancro, IG 13447 and 22221), Wellcome Trust Pathfinder Award (204458/Z/16/Z), Biotechnology and Biological Sciences Research Council (BB/L020874/1 and BB/P018726/1) and CRUK Pioneer Award (29264). MY was supported by the Francis Crick Institute, which receives its core funding from Cancer Research UK (FC001223), the UK Medical Research Council (FC001223) and the Wellcome Trust (FC001223, FC0010060); MY and GS was supported by CRUK Grand Challenge Award 2015 C57633/A25043. MM and RB were supported by the Doctoral School Of Research In Biosciences And Biotechnologies at the University of Padova, and a COMPLeX UCL/British Heart Foundation PhD studentship, respectively. ZR is currently supported by the British Heart Foundation (FS/20/4/34958). RCS was supported by the National Institute for Health Research University College London Hospitals Biomedical Research Centre.

## Author Contributions

Conceptualization: RB, MM, KB, MY, RCS, GS; Methodology: RB, MM, MY, GS; Software: RB, KB, GS; Formal Analysis: RB, MM, NP, SQN, TH, AG, ZR, CE, GS; Investigation: RB, MM, TH, ZR, CE, SA, CL, SI, ZS, MS, AG, MY, GS; Writing – Original Draft: GS; Writing – Review & Editing: RB, MM, MY, GS; Funding Acquisition: MY, GS.

## Declaration of interests

The authors declare no competing interests. RCS is the Chief Investigator of the OPTIMA Trial (ISRCTN42400492) which uses the Prosigna test to make therapeutic decisions in breast cancer treatment. He has no personal financial links with the vendors, NanoString Inc. and Veracyte Inc.

## Notes

**Conflict of interest statement:** The authors declare no competing interests. RCS is the Chief Investigator of the OPTIMA Trial (ISRCTN42400492) which uses the Prosigna test to make therapeutic decisions in breast cancer treatment. He has no personal financial links with the vendors, NanoString Inc. and Veracyte Inc.

### Summary of Updates

Introduction and discussion has been extensively rewritten to reflect and explain better the aims of the analysis. Figures and data remained essentially the same.

