## Supplementary Figures for "Discovery of multi-state gene cluster switches determining the adaptive mitochondrial and metabolic landscape of breast cancer"

### Supplementary Information

**Figure S1.**

**A-C.** Identification and initial characterisation of biclusters from the METABRIC dataset. Silhouette analysis of biclusters identified using the MB1 (**A**) and MB2/MB3 (**B**) genesets (see methods). **C.** Comparison of whole transcriptome CV values between all biclusters found. Density plots of the distribution of CVs across the transcriptome are shown for each bicluster in the central plots. Nuclear encoded mitochondrial genes are shown in red, non-mitochondrial genes are in cyan. The red dashed square indicates the three biclusters identified as independent, and carried over for further analysis.

**D-E.** Identification of the MB1-MB3 biclusters in independent datasets. **D** shows comparison of whole transcriptome CV values from METABRIC, TCGA-microarray, TCGA-RNAseq and Oslo2 datasets. Density plots of the distribution of CVs across the transcriptome are shown for each bicluster/dataset in the central plots. Nuclear encoded mitochondrial genes are shown in red, non-mitochondrial genes are in cyan. **E** shows the MB1 switch in 2D distribution by PC1 and ranking index in the four different dataset. Pam50 classifiers are overlaid to show the similarity of sample distribution in the different switches (biclusters). The result is representative of all three (MB1-MB3 biclusters).

**F**. Average gene group expression values in the upper and lower switch position in the MB1 switch in all datasets (METABRIC, Oslo2, TCGA-microarray, TCGA-RNAseq and TCGA-RRPA). Gene group1 (left panel) and gene group2 (right panel) are the two anticorrelated gene groups discovered by MCbiclust, as shown in Figure 1A. Accordingly, gene group1 shows positive correlation, and higher expression in upper fork samples as compared to the lower fork where conversely, group2 shows positive correlation and higher expression. The direction is arbitrarily set and harmonised between the biclusters. Turkey's box and whisker plots of the mean of the average gene group expression values across the indicated sample group (Lower, Upper Forks). The result is representative of all three (MB1-MB3) biclusters.

**Figure S2.**

**A.** Mapping of highly scoring metabolic pathways from the METABRIC dataset on the global metabolic network. The MB1_UF (top left panel) vs MB1_LF switch (top right panel, blue) is represented. Pathways where any of the enzyme transcripts had a CV over 0.5 (MB1_UF, orange monocolor scale between CV = 0.5-1.0) or under -0.5 (MB1_LF, blue monocolor scale between CV = -0.5 - -1.0) are highlighted. The lower panels show the main domains of the global metabolic network (KEGG, lower left), and the difference between CVs in the two switch positions (lower right, red (max) - white (0) - blue (min) color scale).

**B.** Intra-pathway details of individual gene transcript CVs. Supplementary to Figure 2A, showing pathways with high diversity. CV values of individual genes in the METABRIC breast cancer dataset analysis shown for the KEGG pathways: oxidative phosphorylation, fatty acid synthesis and degradation, purine and pyrimidine synthesis. Values are ranked by the MB_1 CV value. The figure allows the identification of high CV genes and where differences are large between the switches (biclusters).

**Figure S3.**

Statistical analysis of PARADIGM parameters across switch positions. Average PARADIGM values were calculated for each gene in each P cluster (Table 3), and compared between samples of each switch position. Kruskall-Wallis non-parametric test with Holm-corrected pairwise Dunn test was applied.

**Figure S4.**

**A.** Overlap between classification of METABRIC breast cancer samples by switch positions (MB1-3 UF and LF) and PAM50 intrinsic subtypes[57].

**B.** Distribution of different cell-of-origin gene expression scores across the MB1 and MB2 switches.

**Figure S5.**

**A.** Heatmap of fractional enrichment (FE) of $^{U13}$C-glucose derived carbons in isotopologues of key metabolites analysed in MB1_LF (left panel) and MB1_UF (right panel) cells.

**B.** Overall carbon contribution into key metabolites following $^{U13}$C-glucose (top left panel), $^{U13}$C-glutamine (top right panel) and $^{U13}$C-pyruvate (bottom panel) labelling in in MB1_LF and MB1_UF cells under different conditions (see Figure 4A).

**Figure S6.**

**A.** Quantitative analysis of enzyme expression levels obtained from western blot analysis (Fig. 5A**).** Values were normalised to either mitochondrial (left panel) or cytosolic (right panel) markers, grp75 or β-actin. 2-way ANOVA with uncorrected Fisher's LSD multiple comparison tests.

**Figure S7.**

**A.** Visualisation of further histological feature distributions on the MB1 and MB2 switches (see Figure 6B). Classifiers of mitotic activity (left panel), fraction of epithelial area (middle panel) and presence of inflammation (right panel) are overlaid onto 2D distribution plots of TCGA samples along the axes representing the scale between the upper (UF) and lower (LF) forks of each bicluster.

**Tables list S1-8**

**Table S1. METABRIC data: sample characteristics and gene sets**

**Table S2. Unique and overlapping genesets in the MB1 and MB2 switches, together with CV values of the whole transcriptome**

**Table S3. Gene set enrichment analysis (KEGG)**

**Table S4. Gene set enrichment analysis (Mitocarta)**

**Table S5. Top KEGG and Mitocarta pathways**

**Table S6. PARADIGM gene sets**

**Table S7. Breast cancer cell line scoring results**

**Table S8. TCGA BRCA sample classification data with histological and overall features**

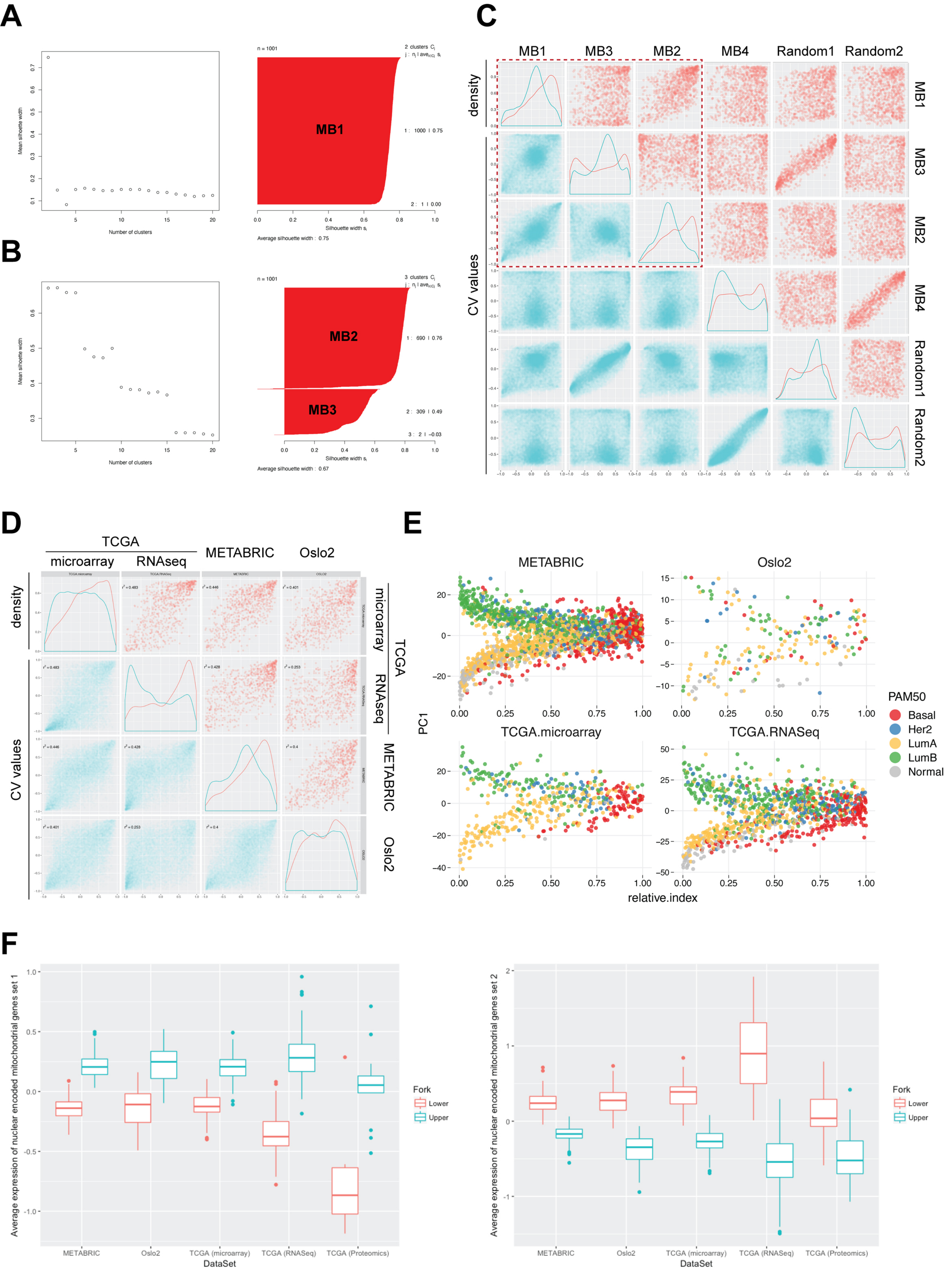

A
Mean silhouette width
Number of clusters
n = 1001
2 clusters
MB1
1 : 1000 | 0.75
2 : 1 | 0.00
Silhouette width
Average silhouette width : 0.75
B
n = 1001
3 clusters
MB2
1 : 690 | 0.76
MB3
2 : 309 | 0.49
3 : 2 | −0.03
Average silhouette width : 0.67
C
MB1
MB3
MB2
MB4
Random1
Random2
density
CV values
D
TCGA
microarray
RNAseq
METABRIC
Oslo2
E
METABRIC
Oslo2
TCGA.microarray
TCGA.RNASeq
PC1
relative.index
PAM50
Basal
Her2
LumA
LumB
Normal
F
Average expression of nuclear encoded mitochondrial genes set 1
Average expression of nuclear encoded mitochondrial genes set 2
Fork
Lower
Upper
METABRIC
Oslo2
TCGA (microarray)
TCGA (RNASeq)
TCGA (Proteomics)
DataSet


### Figure S1

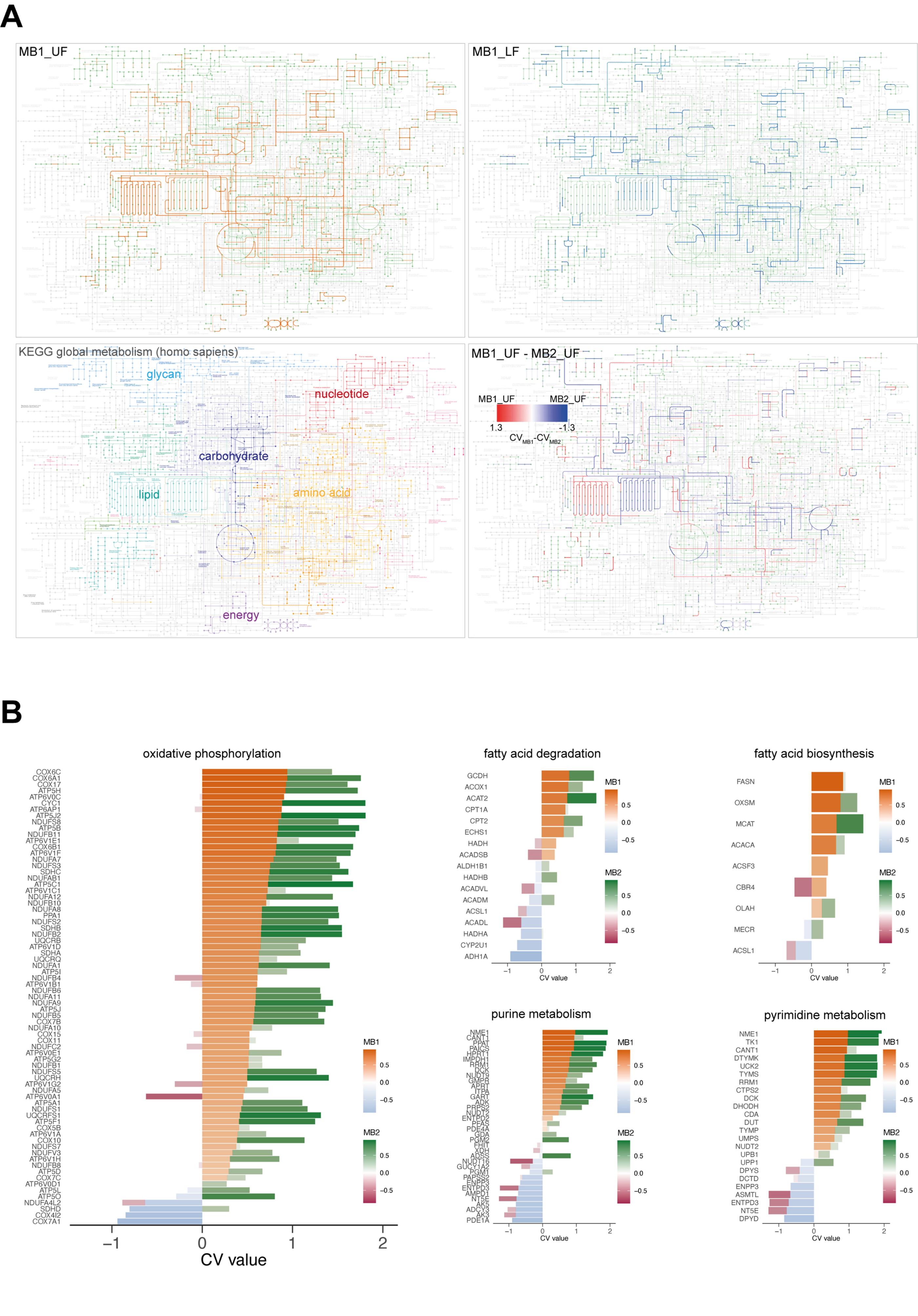
A
MB1_UF
MB1_LF
KEGG global metabolism (homo sapiens)
glycan
nucleotide
carbohydrate
lipid
amino acid
energy
MB1_UF - MB2_UF
MB1_UF
MB2_UF
1.3
-1.3
CV_MB1-CV_MB2
B
oxidative phosphorylation
COX6C
COX6A1
COX17
ATP5H
ATP6V0C
CYC1
ATP6AP1
ATP5J2
NDUFS8
ATP5B
NDUFB11
ATP6V1E1
COX6B1
ATP6V1F
NDUFA7
NDUFS3
SDHC
NDUFAB1
ATP5C1
ATP6V1C1
NDUFA12
NDUFB10
NDUFA8
PPA1
NDUFS2
SDHB
NDUFB2
UQCRB
ATP6V1D
SDHA
UQCRQ
NDUFA1
ATP5I
NDUFB4
ATP6V1B1
NDUFB6
NDUFA11
NDUFA9
ATP5J
NDUFB5
COX7B
NDUFA10
COX15
COX11
NDUFC2
ATP6V0E1
ATP5G2
NDUFB1
NDUFS5
UQCRH
ATP6V1G2
NDUFA5
ATP6V0A1
ATP5A1
NDUFS1
UQCRFS1
ATP5F1
COX5B
ATP6V1A
COX10
NDUFS7
NDUFV3
ATP6V1H
NDUFB8
ATP5D
COX7C
ATP6V0D1
ATP5L
ATP5O
NDUFA4L2
SDHD
COX4I2
COX7A1
−1
0
1
2
CV value
MB1
0.5
0.0
−0.5
MB2
0.5
0.0
−0.5
fatty acid degradation
GCDH
ACOX1
ACAT2
CPT1A
CPT2
ECHS1
HADH
ACADSB
ALDH1B1
HADHB
ACADVL
ACADM
ACSL1
ACADL
HADHA
CYP2U1
ADH1A
−1
0
1
2
CV value
MB1
0.5
0.0
−0.5
MB2
0.5
0.0
−0.5
fatty acid biosynthesis
FASN
OXSM
MCAT
ACACA
ACSF3
CBR4
OLAH
MECR
ACSL1
−1
0
1
2
CV value
MB1
0.5
0.0
−0.5
MB2
0.5
0.0
−0.5
purine metabolism
NME1
CANT1
PPAT
PAICS
HPRT1
IMPDH1
RRM1
DCK
NUDT9
GMPR
APRT
ITPA
GART
ADK
PRPS2
NUDT2
ENTPD2
PFAS
PDE4A
GDA
PGM2
FHIT
XDH
ADSS
NUDT16
GUCY1A2
PGM1
PAPSS2
ENPP3
ENTPD3
AMPD1
NT5E
AK5
ADCY3
AK3
PDE1A
−1
0
1
2
CV value
MB1
0.5
0.0
−0.5
MB2
0.5
0.0
−0.5
pyrimidine metabolism
NME1
TK1
CANT1
DTYMK
UCK2
TYMS
RRM1
CTPS2
DCK
DHODH
CDA
DUT
TYMP
UMPS
NUDT2
UPB1
UPP1
DPYS
DCTD
ENPP3
ASMTL
ENTPD3
NT5E
DPYD
−1
0
1
2
CV value
MB1
0.5
0.0
−0.5
MB2
0.5
0.0
−0.5

### Figure S2

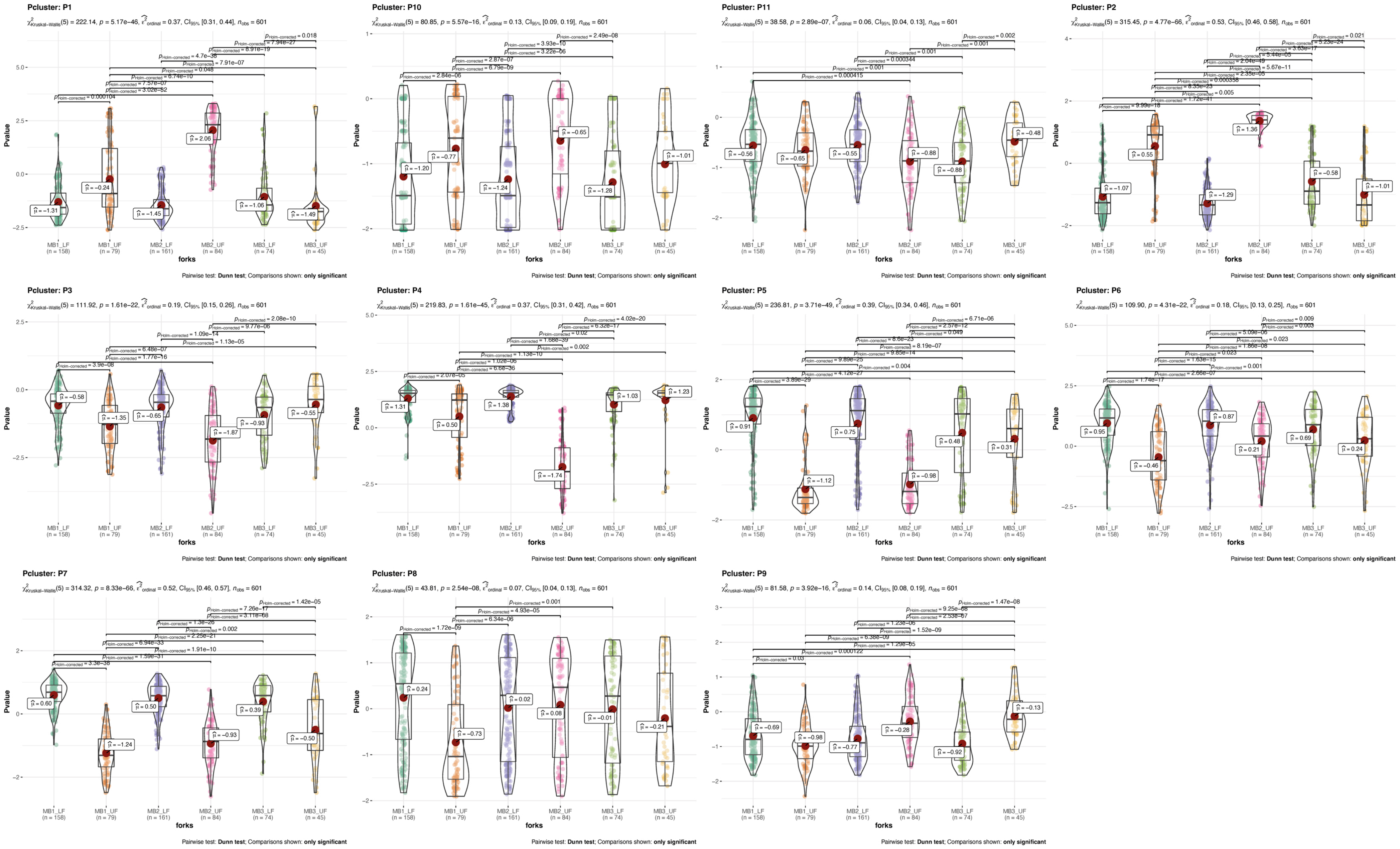

Pcluster: P1
χ²Kruskal-Wallis(5) = 222.14, p = 5.17e-46, ε²ordinal = 0.37, CI95% [0.31, 0.44], nobs = 601
Pcluster: P10
χ²Kruskal-Wallis(5) = 80.85, p = 5.57e-16, ε²ordinal = 0.13, CI95% [0.09, 0.19], nobs = 601
Pcluster: P11
χ²Kruskal-Wallis(5) = 38.58, p = 2.89e-07, ε²ordinal = 0.06, CI95% [0.04, 0.13], nobs = 601
Pcluster: P2
χ²Kruskal-Wallis(5) = 315.45, p = 4.77e-66, ε²ordinal = 0.53, CI95% [0.46, 0.58], nobs = 601
Pcluster: P3
χ²Kruskal-Wallis(5) = 111.92, p = 1.61e-22, ε²ordinal = 0.19, CI95% [0.15, 0.26], nobs = 601
Pcluster: P4
χ²Kruskal-Wallis(5) = 219.83, p = 1.61e-45, ε²ordinal = 0.37, CI95% [0.31, 0.42], nobs = 601
Pcluster: P5
χ²Kruskal-Wallis(5) = 236.81, p = 3.71e-49, ε²ordinal = 0.39, CI95% [0.34, 0.46], nobs = 601
Pcluster: P6
χ²Kruskal-Wallis(5) = 109.90, p = 4.31e-22, ε²ordinal = 0.18, CI95% [0.13, 0.25], nobs = 601
Pcluster: P7
χ²Kruskal-Wallis(5) = 314.32, p = 8.33e-66, ε²ordinal = 0.52, CI95% [0.46, 0.57], nobs = 601
Pcluster: P8
χ²Kruskal-Wallis(5) = 43.81, p = 2.54e-08, ε²ordinal = 0.07, CI95% [0.04, 0.13], nobs = 601
Pcluster: P9
χ²Kruskal-Wallis(5) = 81.58, p = 3.92e-16, ε²ordinal = 0.14, CI95% [0.08, 0.19], nobs = 601
Pvalue
forks
MB1_LF (n = 158)
MB1_UF (n = 79)
MB2_LF (n = 161)
MB2_UF (n = 84)
MB3_LF (n = 74)
MB3_UF (n = 45)
Pairwise test: Dunn test; Comparisons shown: only significant


Figure S3

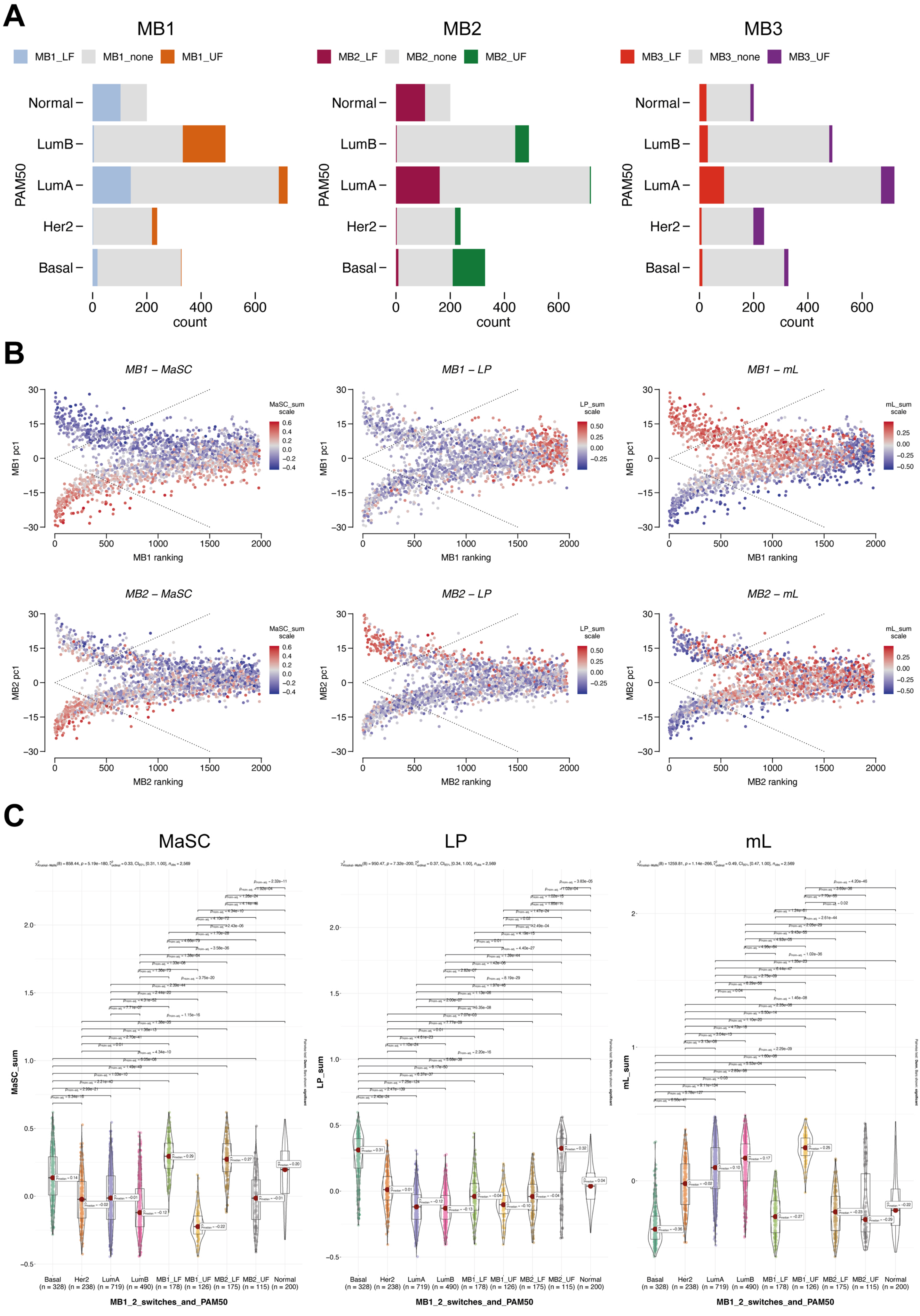
A
MB1
MB1_LF
MB1_none
MB1_UF
MB2
MB2_LF
MB2_none
MB2_UF
MB3
MB3_LF
MB3_none
MB3_UF
PAM50
Normal
LumB
LumA
Her2
Basal
0
200
400
600
count
B
MB1 – MaSC
MB1 – LP
MB1 – mL
MB2 – MaSC
MB2 – LP
MB2 – mL
MB1 pc1
MB2 pc1
MB1 ranking
MB2 ranking
MaSC_sum scale
LP_sum scale
mL_sum scale
C
MaSC
LP
mL
MaSC_sum
LP_sum
mL_sum
Basal (n = 328)
Her2 (n = 238)
LumA (n = 719)
LumB (n = 490)
MB1_LF (n = 178)
MB1_UF (n = 126)
MB2_LF (n = 175)
MB2_UF (n = 115)
Normal (n = 200)
MB1_2_switches_and_PAM50

Figure S4

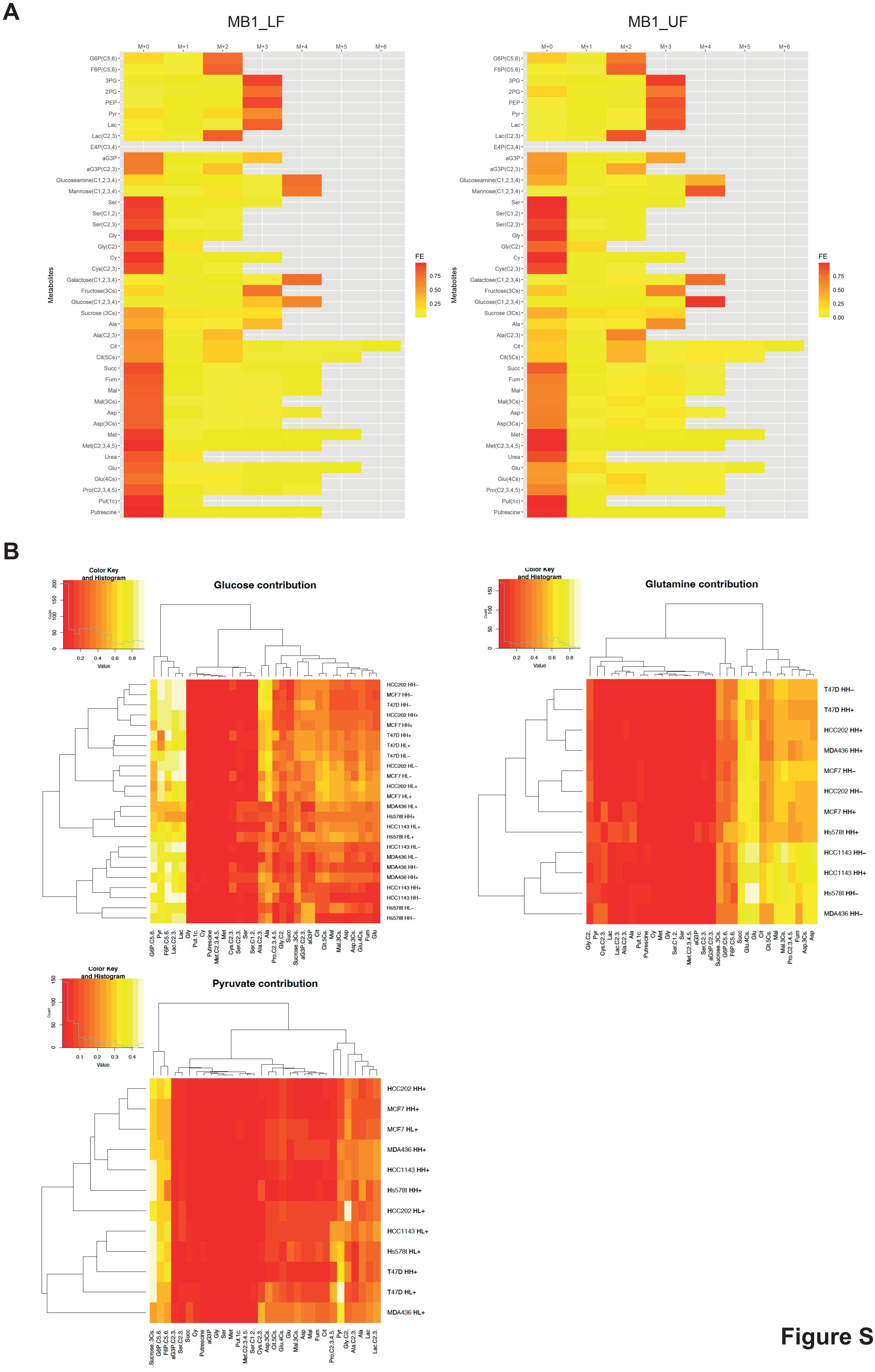
A
MB1_LF
MB1_UF
M+0
M+1
M+2
M+3
M+4
M+5
M+6
G6P(C5,6)
F6P(C5,6)
3PG
2PG
PEP
Pyr
Lac
Lac(C2,3)
E4P(C3,4)
aG3P
aG3P(C2,3)
Glucoseamine(C1,2,3,4)
Mannose(C1,2,3,4)
Ser
Ser(C1,2)
Ser(C2,3)
Gly
Gly(C2)
Cy
Cys(C2,3)
Galactose(C1,2,3,4)
Fructose(3Cs)
Glucose(C1,2,3,4)
Sucrose (3Cs)
Ala
Ala(C2,3)
Cit
Cit(5Cs)
Succ
Fum
Mal
Mal(3Cs)
Asp
Asp(3Cs)
Met
Met(C2,3,4,5)
Urea
Glu
Glu(4Cs)
Pro(C2,3,4,5)
Put(1c)
Putrescine
Metabolites
FE
0.75
0.50
0.25
0.00
B
Color Key and Histogram
Count
Value
Glucose contribution
Glutamine contribution
Pyruvate contribution
HCC202 HH–
MCF7 HH–
T47D HH–
HCC202 HH+
MCF7 HH+
T47D HH+
T47D HL+
T47D HL–
HCC202 HL–
MCF7 HL–
HCC202 HL+
MCF7 HL+
MDA436 HL+
Hs578t HH+
HCC1143 HL+
Hs578t HL+
HCC1143 HL–
MDA436 HL–
MDA436 HH–
MDA436 HH+
HCC1143 HH+
HCC1143 HH–
Hs578t HL–
Hs578t HH–
Hs578t HH–
HCC202 HL+
HCC1143 HL+
Hs578t HL+
T47D HL+
MDA436 HL+

Figure S5

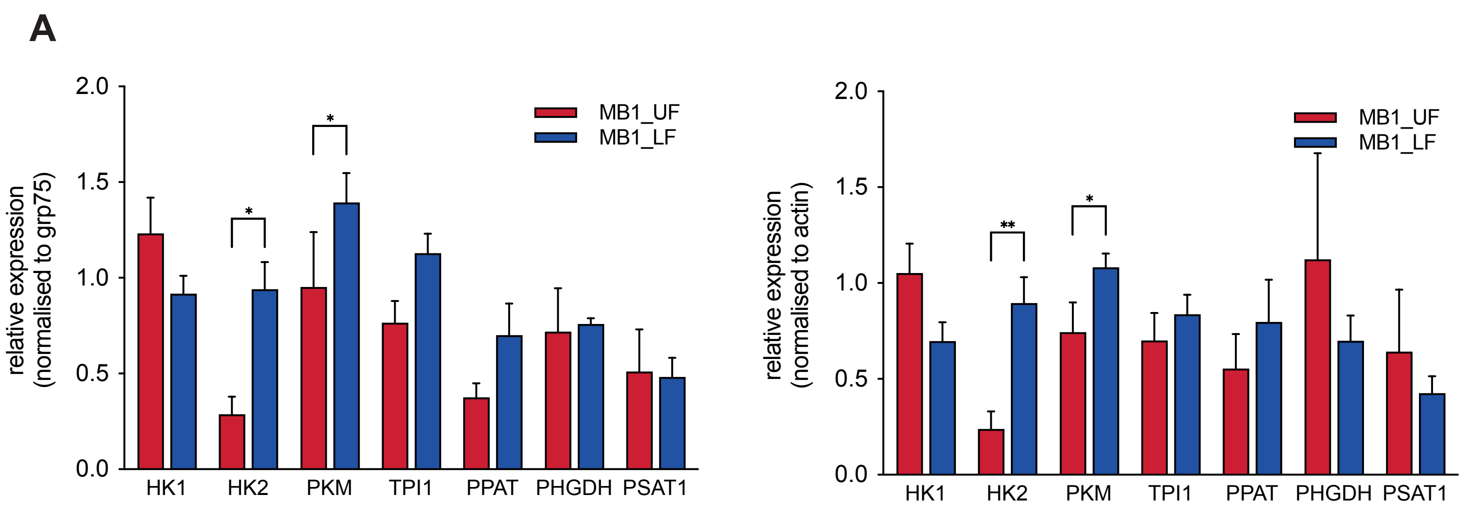
A
MB1_UF
MB1_LF
relative expression (normalised to grp75)
relative expression (normalised to actin)
2.0
1.5
1.0
0.5
0.0
*
**
HK1
HK2
PKM
TPI1
PPAT
PHGDH
PSAT1

Figure S6

A

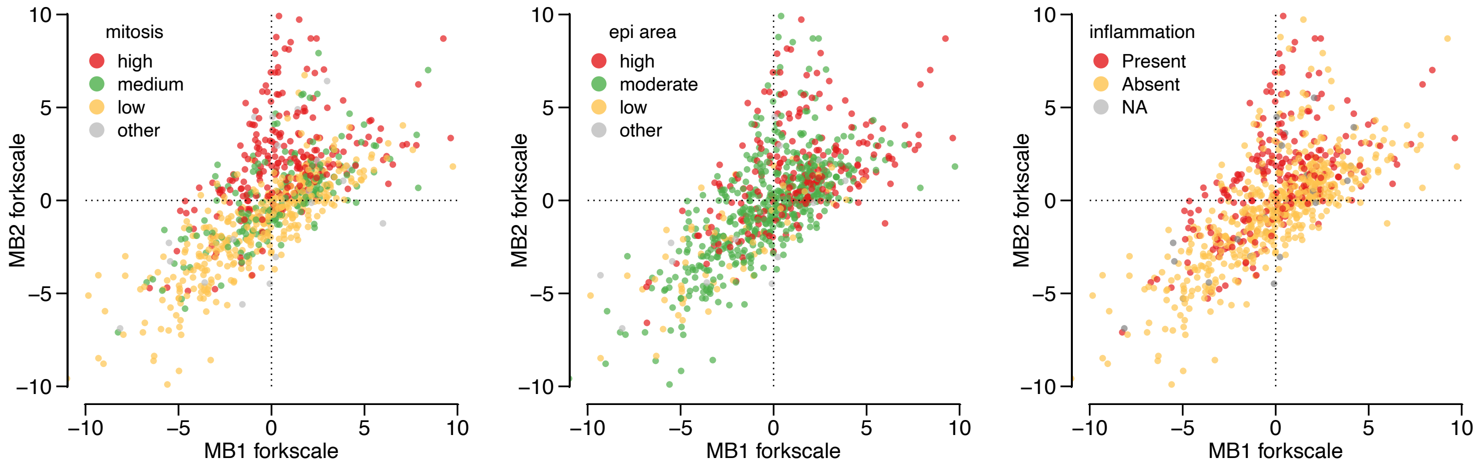

mitosis
high
medium
low
other
MB2 forkscale
MB1 forkscale
epi area
high
moderate
low
other
MB2 forkscale
MB1 forkscale
inflammation
Present
Absent
NA
MB2 forkscale
MB1 forkscale


Figure S7
